# Piezo2 in sensory neurons influences systemic and adipose tissue metabolism

**DOI:** 10.1101/2024.11.27.624676

**Authors:** Fabian S. Passini, Bavat Bornstein, Sarah Rubin, Yael Kuperman, Sharon Krief, Evi Masschelein, Tevie Mehlman, Alexander Brandis, Yoseph Addadi, Shira Huri-Ohev Shalom, Erik A. Richter, Tal Yardeni, Amir Tirosh, Katrien De Bock, Elazar Zelzer

## Abstract

Systemic metabolism ensures energy homeostasis through inter-organ crosstalk regulating thermogenic adipose tissue. Unlike the well-described inductive role of the sympathetic system, the inhibitory signal ensuring energy preservation remains poorly understood. Here, we show that, via the mechanosensor Piezo2, sensory neurons regulate morphological and physiological properties of brown and beige fat and prevent systemic hypermetabolism. Targeting Runx3/PV sensory neurons in independent genetic mouse models resulted in a systemic metabolic phenotype characterized by reduced body fat and increased insulin sensitivity and glucose tolerance. Deletion of Piezo2 in PV sensory neurons reproduced the phenotype, protected against high-fat diet-induced obesity and caused adipose tissue browning and beiging, likely driven by elevated norepinephrine levels. Finding that brown and beige fat are innervated by Runx3/PV sensory neurons expressing Piezo2, suggests a model where mechanical signals sensed by Piezo2 in sensory neurons protect energy storages and prevent a systemic metabolic phenotype.

**Highlights:** Lack of Runx3/PV sensory neurons reduces body fat and fasting glucose levels and increases glucose tolerance in mice

Mechanosensitive ion channel PIEZO2 in PV sensory neurons plays an important role in systemic metabolism under physiological and pathological conditions

PIEZO2 in PV sensory neurons regulates thermogenic programs and glucose uptake in brown and beige adipose tissues

Brown and beige adipose tissues are innervated by Runx3/PV sensory neurons

## Introduction

Systemic metabolism has the remarkable ability to adapt to diverse challenges, enabling organisms to thrive and survive in varying conditions (*1, 2*). The impact of a dysfunctional systemic metabolism has become increasingly evident in recent decades with the substantial increase in the prevalence of obesity and type 2 diabetes (*3*).

Adipose tissue plays a pivotal role in regulating systemic metabolism. Beyond its fundamental function of storing triglycerides as an energy reserve and releasing fatty acids for other tissues, it affects key aspects of whole-body energy homeostasis. These include the regulation of food intake, glucose homeostasis, insulin sensitivity, and thermogenesis (*1, 4, 5*). Adipose tissue is broadly categorized into two forms: white adipose tissue (WAT) and brown adipose tissue (BAT). White adipocytes are characterized by numerous unilocular lipid droplets and limited mitochondria, whereas brown adipocytes feature small multilocular lipid droplets and abundant mitochondria, which are essential for their specialized role in heat production (*6*). Thermogenesis results from the uncoupling of electron transfer through the respiratory chain from ATP production, mediated by the activity of uncoupling protein-1 (UCP-1) as well as by UCP-1 independent mechanisms (*6–8*). The thermogenic activity of BAT significantly enhances the uptake of fatty acids and glucose as energy substrates, contributing substantially to whole-body glucose disposal and energy expenditure (*9*). However, thermogenic capacity is not exclusive to brown adipocytes. Rather, it can also be observed in brown-like adipocytes within white adipose tissue, a phenomenon referred to as browning or beiging (*10*). These beige adipocytes share characteristics with brown adipocytes, such as multilocular lipid droplets and denser mitochondria. The browning process is regulated by various stimuli, including cold exposure, catecholamines, exercise, thiazolidinediones, and injury (*10, 11*).

Adipose tissues are innervated by both efferent sympathetic and afferent sensory neurons, forming a feedback loop that allows bidirectional communication between fat and the central nervous system (*12–14*). Previous studies have elucidated the central role of sympathetic innervation in regulating various aspects of adipose tissues, including thermogenesis, lipolysis, and adipogenesis (*15*). These processes are mediated by the release of norepinephrine from sympathetic nerve terminals, activating thermogenic programs in both brown and beiging adipocytes expressing its β3-adrenergic receptor (*12, 15*).

Although adipose sensory innervation has long been known, our understanding of its function lags behind that of the sympathetic system. Recent tissue ablation experiments targeting the sensory system locally have demonstrated its involvement in regulating the browning of white fat (*16*). However, gaps persist in our understanding of the signals sensed by these sensory nerves, the global effects of the sensory system on adipose tissue, and its role in regulating systemic metabolism. The unknown identity of adipose sensory neurons has hindered this exploration.

In this study, we discovered an important role of the sensory system and the mechanosensitive ion channel Piezo2 in regulating brown and beige fat properties and systemic metabolism in both physiological and pathological conditions.

## Results

### Deletion of Runx3/PV sensory neurons leads to a systemic metabolic phenotype

While investigating the role of sensory neurons in musculoskeletal physiology (*17*), we noticed that mice lacking expression of the runt-domain transcription factor 3 (Runx3) had reduced fat depots. Runx3 is necessary for the development and survival of TrkC sensory neurons (*18*). To investigate in detail this initial observation, we assessed total body fat of *Runx3* knockout (KO) mice using a body composition analysis. *Runx3* KO mice showed reduced body weight with decreased body fat percentage (Fig. 1a and Fig. S1a). As this reduction could result from abnormal systemic metabolism (*4*), we proceeded with metabolic profiling using metabolic cages. Examination of respiratory exchange ratio (RER), the ratio between carbon dioxide release and oxygen consumption, revealed a stronger oscillation in *Runx3* KO mice (Fig. 1b). This is likely a result of differences in energy balance between mutants and controls (*19*). *Runx3* KO mice also exhibited reduced mobility (Fig. 1b, Fig. S1). Next, we assessed glucose metabolism and insulin sensitivity in these mice. Importantly, we found reduced fasting blood glucose levels and improved glucose tolerance in *Runx3* KO mice (Fig. 1c,d). In addition, these mice displayed reduced levels of insulin and increased insulin sensitivity (Fig. 1e,f). Hence, our analyses revealed a strong metabolic phenotype characterized by excessive glucose uptake, reduced fasting glucose levels and decreased body fat percentage in *Runx3* KO mice.

**Fig. 1.**
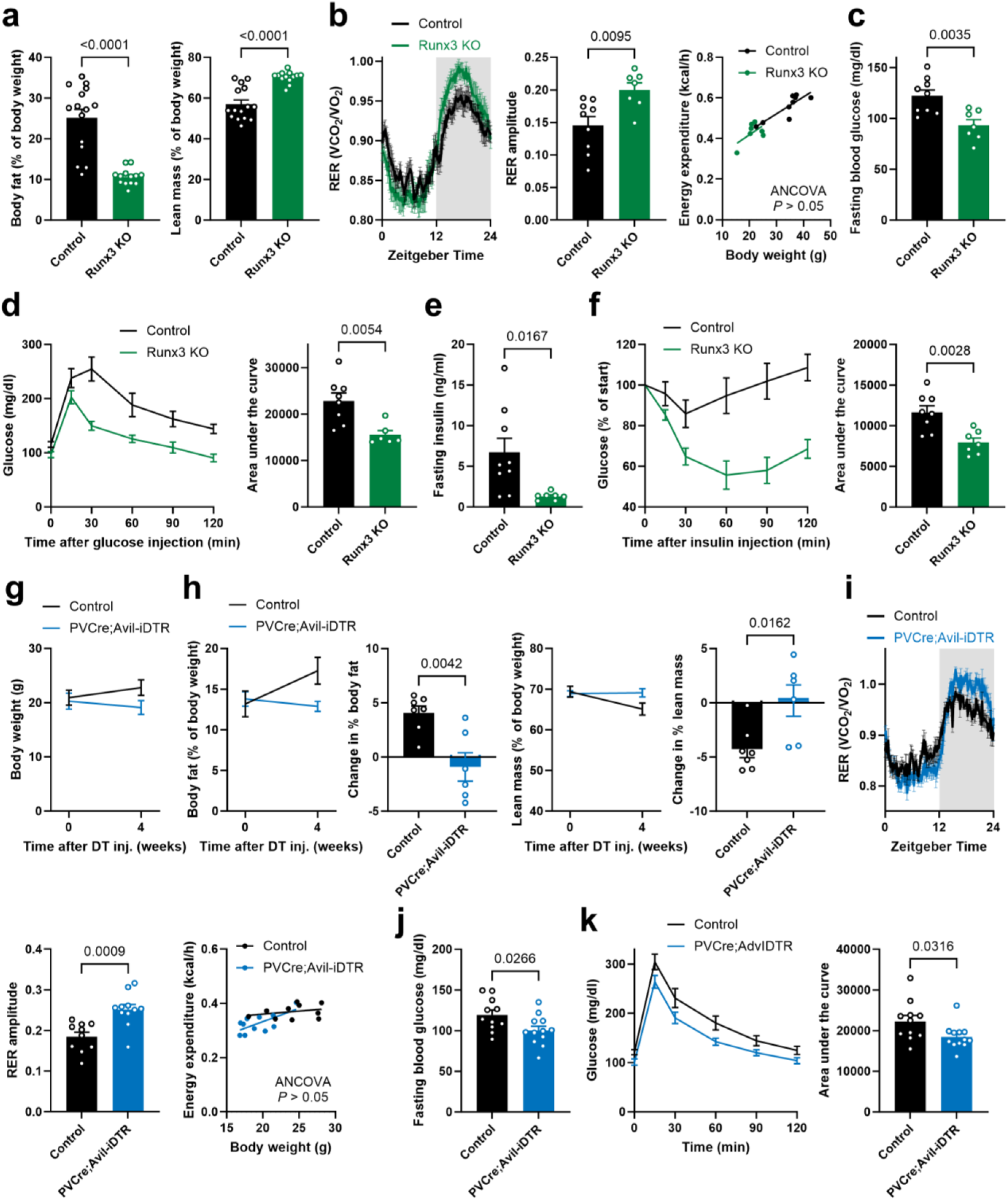
*Runx3* KO mice (ICR background) and *PV-Cre;Avil-iDTR* mice (C57BL/6 background) show a systemic metabolic phenotype. a) Body composition analysis revealed reduced body fat and increased lean mass as a percentage of body weight in *Runx3* KO mice (n = 13) relative to littermate controls (n = 15). b-f) Graphs showing metabolic changes in *Runx3* KO mice (n = 6-7) as compared to control littermates (n = 8-9). b) Metabolic cage analysis showed a stronger RER oscillation between the light (0-12 h) and dark phase (12-24 h) in *Runx3* KO mice, while energy expenditure remained comparable. c) Reduced blood glucose after 6 h fasting. d) Glucose tolerance test (2 g/kg body weight) performed after 6 h fasting showed improved glucose uptake. e) Reduced plasma insulin level after 6 h fasting. f) Insulin tolerance test (0.6 U/kg body weight) performed after 3 h fasting shows increased insulin sensitivity. *Runx3 KO* mice and littermate controls were 2-5 months old at the timepoint of the experiments. g, h) Diphtheria toxin was injected into 2-3 months old *PV-Cre;Avil-iDTR* mice (n = 6) and control littermates (n = 7). Body weight (g) and body composition (h) were analyzed before and 4 weeks after the injection. Changes in body fat and lean mass as a percentage of body weight were observed 4 weeks after diphtheria toxin injection. i-k) Graphs showing metabolic changes in *PV- Cre;Avil-iDTR* mice (n = 11-12) as compared to control littermates (n = 10-11). i) Metabolic cage analysis showed a stronger RER oscillation in *PV-Cre;Avil-iDTR* mice, while energy expenditure remained comparable. j) Reduced blood glucose after 6 h fasting. k) Glucose tolerance test (2 g/kg body weight) performed after 6 h fasting showed improved glucose uptake. *PV-Cre;Avil-iDTR* mice and littermate controls were 2-3 months old at the time of diphtheria toxin injection and metabolic tests were performed 1-2 months later. Mice were kept at room temperature of around 22°C. Unless indicated otherwise, statistical analysis was performed using unpaired t- tests. ANCOVA analysis was performed with body weight as covariate. Data are presented as mean ± SEM.

The vast majority of Runx3 sensory neurons express parvalbumin (PV) (*20*). To test whether the observed metabolic phenotype in *Runx3* KO mice is potentially caused by sensory neurons, we specifically ablated PV sensory neurons in adult mice by crossing *PV-Cre* mice with *Avil-iDTR* mice, where expression of the human diphtheria toxin receptor (DTR) in peripheral sensory neurons is driven by the pan-somatosensory marker advillin (*21–23*). Four weeks after the injection of diphtheria toxin, we found that control mice gained body weight, whereas *PV-Cre;Avil-iDTR* mice exhibited a slight decrease (Fig. 1g). This difference is primarily due to changes in body fat that increased as a percentage of body weight in controls but decreased in *PV-Cre;Avil-iDTR* mice (Fig. 1h and Fig. S2a). Consequently, the lean mass percentage decreased in the controls but not in *PV-Cre;Avil-iDTR* mice (Fig. 1h). Metabolic tests showed that, similar to *Runx3* KO mice, *PV-Cre;Avil-iDTR* mice displayed stronger RER oscillations, reduced fasting blood glucose levels and increased glucose tolerance, along with an unchanged food intake (Fig. 1i-k, Fig. S2b-g). These findings suggest an important regulatory role of Runx3/PV sensory neurons in maintaining metabolic homeostasis and preserving body fat storages and blood glucose levels.

### Deletion of the mechanosensor Piezo2 in PV sensory neurons reproduces systemic metabolic phenotype

PV sensory neurons were shown to express Piezo2 (*24*), raising the possibility that this mechanosensitive ion channel is involved in maintaining systemic metabolic homeostasis. Piezo2 detects mechanical forces (*25, 26*) and plays key physiological functions in various subpopulations of sensory neurons. For instance, in interoceptive sensory neurons, Piezo2 detects lung inflation (*27*) and bladder filling (*28*), and in proprioceptive sensory neurons it detects muscle stretch (*24*).

We next examined whether Piezo2 is involved in regulating systemic metabolism by conducting comprehensive metabolic tests on *Piezo2* cKO mice, which lack *Piezo2* expression in PV sensory neurons. Strikingly, we found a clear metabolic phenotype that was consistent with that found in *Runx3* KO and *PV-Cre;Avil-iDTR* mice. *Piezo2* cKO mice displayed lower body weight, primarily due to a reduction in fat mass as observed by the reduced body fat percentage and increased lean mass percentage (Fig. 2a and Fig. S3a). Moreover, we observed an amplified RER oscillation in *Piezo2* cKO mice, along with reduced mobility and similar food intake (Fig. 2b and Fig. S3). *Piezo2* cKO mice also exhibited reduced fasting blood glucose levels and increased glucose tolerance (Fig. 2c,d), as well as reduced fasting insulin levels and increased insulin sensitivity (Fig. 2e,f and Fig. S4). To control for the specificity of the effects observed by blocking *Piezo2* expression in PV sensory neurons, we deleted *Piezo2* using *Scn10a*-driven Cre, a common approach to target a broad range of high-threshold mechanosensory neurons without affecting the PV-population (*29, 30*). This deletion did not lead to any significant changes in body composition, fasting blood glucose levels, glucose tolerance, or insulin sensitivity (Fig. 2g-i and Fig. S6). Finally, we were interested in determining whether the observed effects were temperature-dependent. To test this, we housed *Piezo2* cKO and control mice at a murine thermoneutrality of 30°C from birth (*31*). Under these conditions, *Piezo2* cKO mice exhibited reduced body weight and body fat percentage, while their fasting glucose levels and glucose tolerance remained unaffected (Fig. S5). These results suggest that the metabolic phenotype of *Piezo2* cKO mice is, at least in part, influenced by temperature.

**Fig. 2.**
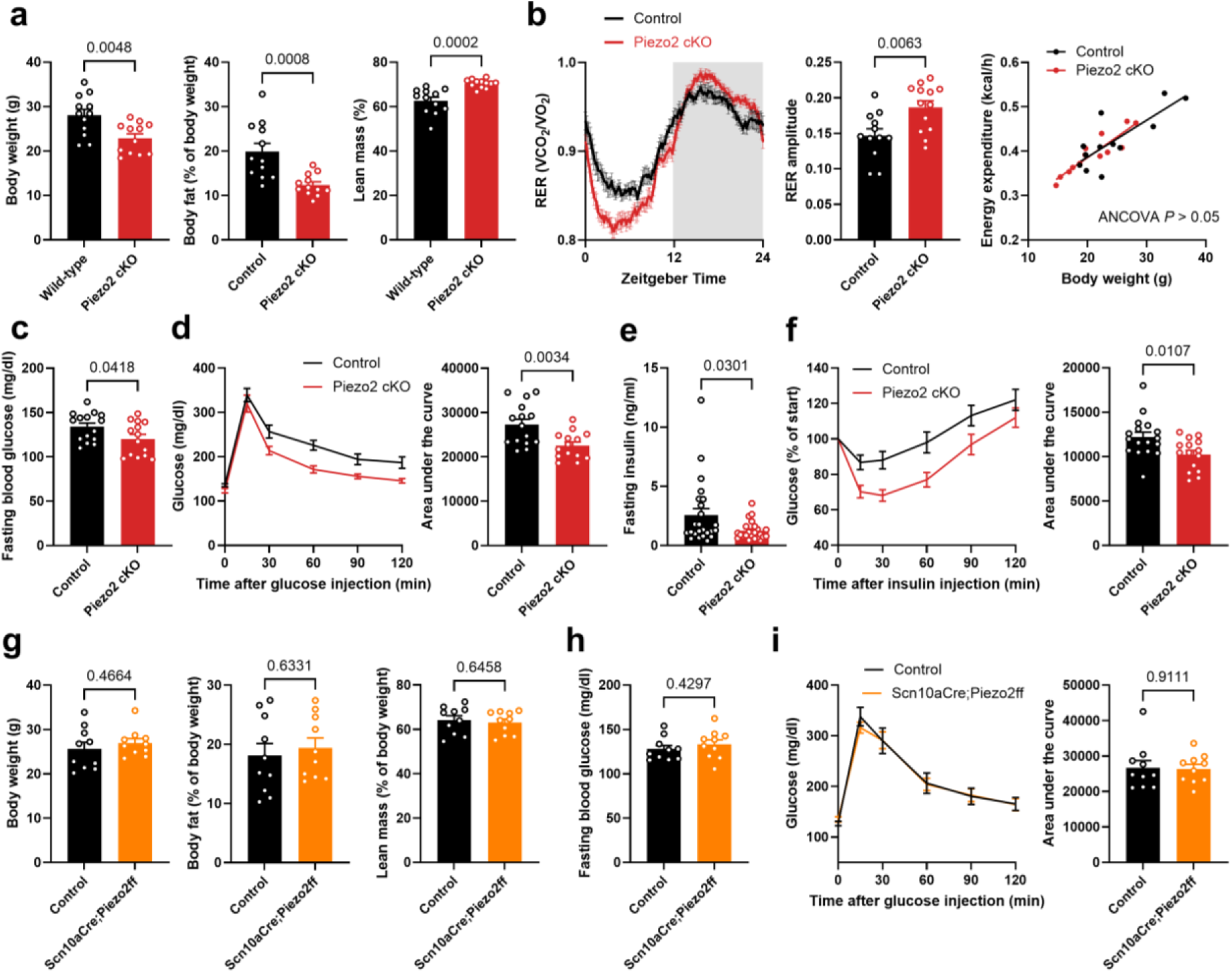
*Piezo2* cKO mice (*PV-Cre;Piezo2flox/flox*, C57BL/6 background) show a systemic metabolic phenotype, while *Scn10a-Cre;Piezo2flox/flox* mice do not exhibit alterations in body composition and glucose metabolism. a) Measurements of body weight and body composition revealed reduced body fat and increased lean mass as a percentage of body weight in *Piezo2* cKO mice (n=12) relative to littermate controls (n = 12). b) Metabolic cage analysis found a stronger RER oscillation in *Piezo2* cKO mice (n=13) relative to littermate controls (n = 12), while energy expenditure remained comparable. ANCOVA analysis was performed with body weight as covariate. c, d) Glucose metabolism analyses performed after 6 h fasting in *Piezo2* cKO mice (n = 14) and littermate controls (n = 15) revealed reduced blood glucose (c) and improved glucose uptake (glucose tolerance test, 2 g/kg body weight; d) in the mutants. e) Reduced plasma insulin level after 6 h fasting in *Piezo2* cKO mice (n = 23) as compared to littermate controls (n = 24). f) Insulin tolerance test (0.6 U/kg body weight) performed after 3 h fasting shows increased insulin sensitivity in *Piezo2* cKO mice (n = 15) as compared to littermate controls (n = 17). g) Unchanged body weight and body composition in *Scn10a- Cre;Piezo2flox/flox* mice (n = 10) compared to littermate controls (n = 10). h, i) Glucose metabolism analyses performed after 6 h fasting in *Scn10a-Cre;Piezo2flox/flox* mice (n = 10) and littermate controls (n = 10) showed unaffected blood glucose (h) and glucose uptake (glucose tolerance test, 2 g/kg body weight; i) in the mutants. Mice were 3-6 months old at the timepoint of the experiments. Mice were kept at room temperature of around 22°C. Unless indicated otherwise, statistical analysis was performed using unpaired t-tests. Data are presented as mean ± SEM.

Taken together, similar metabolic phenotypes were observed upon ablation of Runx3/PV sensory neurons and upon deletion of *Piezo2* in these neurons, but not upon its deletion in the *Scn10a*- positive population. This indicates a key role of Piezo2 within the Runx3/PV sensory population in maintaining energy storages and regulating systemic metabolism.

### Piezo2 deletion in PV sensory neurons protects against high-fat diet-induced metabolic diseases

We next asked whether this metabolic phenotype is protective against metabolic diseases. To address this, we used a diet-induced obesity approach (*32*). We switched the diet of adult *Piezo2* cKO and control mice from a regular chow-diet to a high-fat diet (HFD; 60% of calories from fat) and monitored body weight and body composition during a 3-month period. We observed that the difference in body fat between *Piezo2* cKO and control mice was amplified by the HFD intervention, as the mutants accumulated less body fat (Fig. 3a,b). This finding prompted us to analyze the liver of these animals, given that obesity is often accompanied by hepato-steatosis (*33*). Importantly, we found that liver steatosis was strongly attenuated in *Piezo2* cKO mice relative to control mice, as lipid droplet accumulation in the liver was practically prevented (Fig. 3c,d). Additionally, *Piezo2* cKO mice maintained an improved glucose tolerance and insulin sensitivity during HFD compared with control mice (Fig. 3e,f). These data indicate that the fmetabolic phenotype of *Piezo2* cKO is protective of obesity-related metabolic abnormalities.

**Fig. 3.**
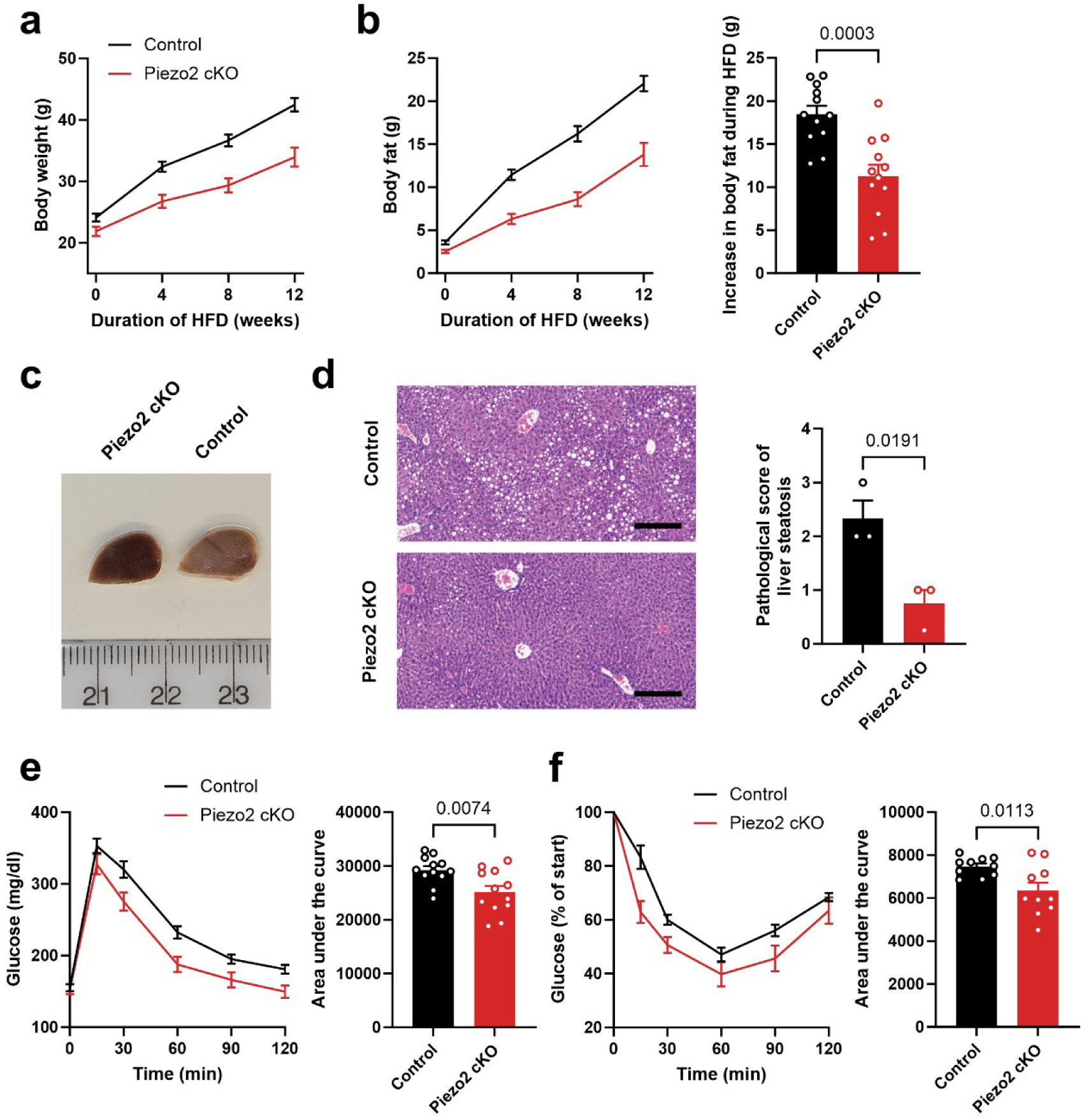
*Piezo2* cKO mice are protected against HFD-induced metabolic disorders, including obesity and liver steatosis. a, b) Graphs showing body weight (a) and body fat (b) during 12 weeks of HFD in *Piezo2* cKO mice and littermate controls (n = 12 for both). c) Image of livers after 12 weeks of HFD. d) H&E staining of the liver after the HFD period. Right: Corresponding pathological scoring for liver steatosis on a scale ranging from 0 (normal) to 4 (severe). Scale bar, 200 µm; n = 3 for both mutants and littermate controls. e,f) Graphs showing improved glucose uptake (glucose tolerance test after 6 h fasting, 1 g/kg body weight; n=12 for both; e) and increased insulin sensitivity (insulin tolerance test after 6 h fasting, 1 U/kg body weight; n=10 for both; f) in *Piezo2* cKO mice relative to littermate controls. Mice were kept at room temperature of around 22°C. Statistical analysis was performed using unpaired t-tests. Data are presented as mean ± SEM.

### Piezo2 deletion in PV sensory neurons alters adipose tissue metabolism but not muscle metabolism

PV-positive sensory neurons are well-known to innervate proprioceptors, such as the muscle spindle (*20, 34*). Moreover, it was shown that *Piezo2* is necessary for proprioception, and that *Piezo2* cKO mice have proprioceptive deficits and exhibit ataxia (*24*). Given that muscles play a key role in systemic metabolism (*2, 35*), we investigated whether they might contribute to the metabolic phenotype of *Piezo2* cKO mice. Thorough analysis of mutant and control mice revealed no significant differences in muscle weight, slightly reduced fiber size but no changes in fiber type composition (Fig. 4a-c). Moreover, comparative transcriptional profiling by RNA sequencing showed that solely 0.29% and 0.04% of all identified genes were differentially expressed in the soleus and gastrocnemius muscles, respectively (Fig. 4d), none of which is known to be involved in key metabolic processes. Thus, muscles in *Piezo2* cKO mice remain largely unaffected.

**Fig. 4.**
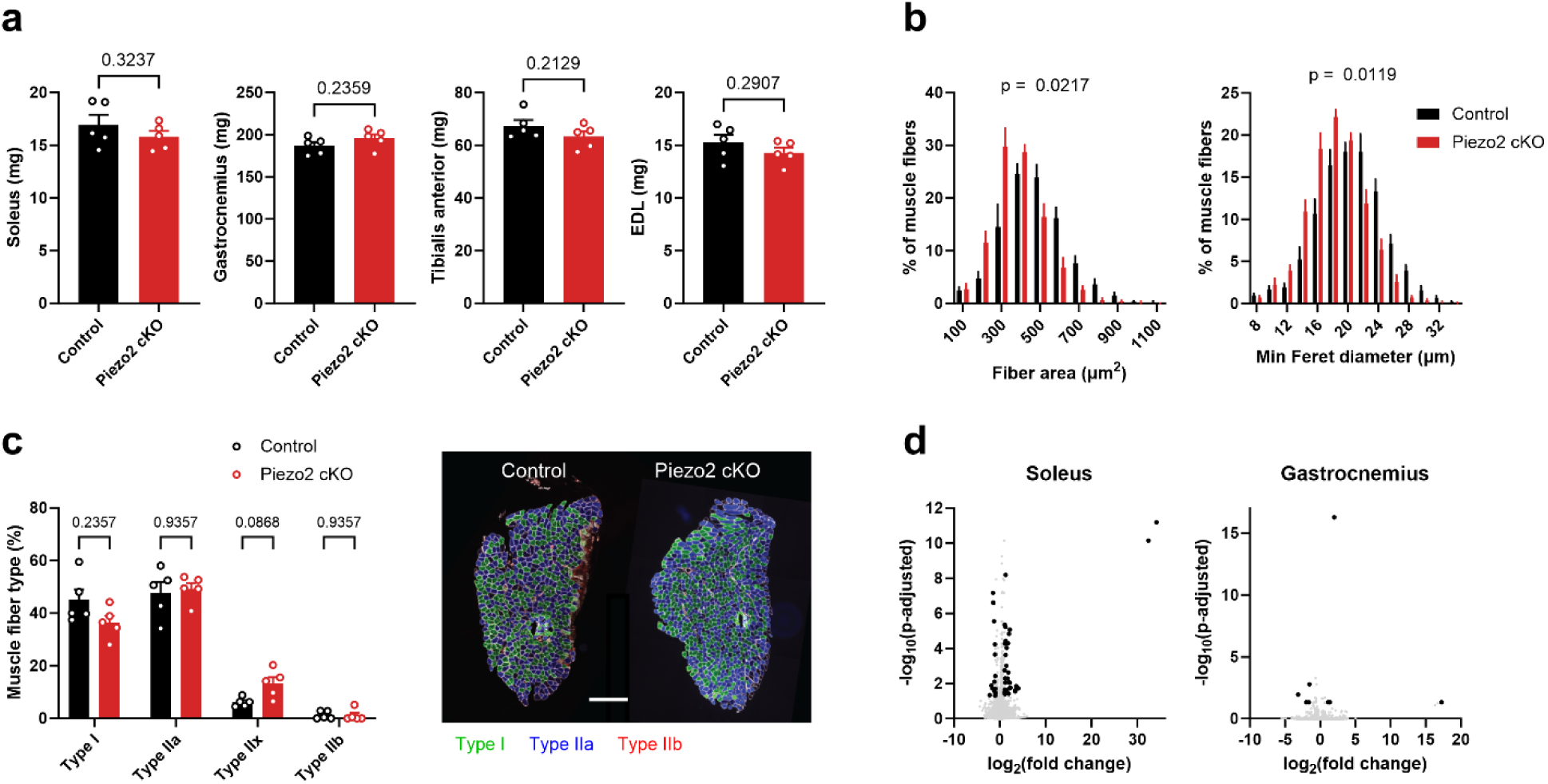
Muscles from *Piezo2* cKO mice show similar weight, slightly reduced fiber size and highly comparable fiber type composition and transcriptional profile. a) Wet weight of soleus, gastrocnemius, tibialis anterior and extensor digitorum longus (EDL) muscles. Statistical analysis was performed using unpaired t-tests. b) Area and min Feret diameter were found to be mildly reduced in the soleus muscle from *Piezo2* cKO mice compared to controls. Statistical analysis was performed using a linear mixed model with the mouse ID as a random factor. c) Fiber type composition of the soleus muscle assessed with immunofluorescence staining, scalebar 200 µm. Statistical analysis was performed using multiple t-tests. d) Transcriptome of soleus and gastrocnemius muscles from bulk RNA-sequencing analysis shown as Volcano plots. Differentially expressed genes (log2 fold change >1, adjusted p-value < 0.05) are labelled in black. In the soleus muscle from Piezo2 cKO mice, 37 genes were found to be upregulated and 16 downregulated compared to control, while in the gastrocnemius muscles 4 genes were upregulated and 4 downregulated. n=5 *Piezo2* cKO mice, n=5 littermate controls. Data are mean ± SEM.

The observed reduction in body fat and increase in glucose tolerance and insulin sensitivity in *Piezo2* cKO mice suggests an elevated utilization of fuels, such as glucose. To identify the tissue(s) where this increased utilization occurs, we took an unbiased approach and examined the *in vivo* glucose uptake by various tissues 30 minutes after the combined injection of radiolabeled 2-deoxyglucose (2-DG) and insulin (Fig. 5a). 2-DG tracing showed increased glucose uptake only in interscapular BAT (iBAT) and inguinal WAT (iWAT, a beige fat pad (*5*)) of *Piezo2* cKO mice relative to the controls, while in other tissues, such as the soleus, gastrocnemius, tibialis anterior, and extensor digitorum longus muscles, visceral fat, heart, liver, or brain, the uptake remained comparable between mutants and controls (Fig. 5b and Fig. S7). These results suggest that Piezo2 in sensory neurons regulates glucose uptake specifically in brown and beige fat.

**Fig. 5.**
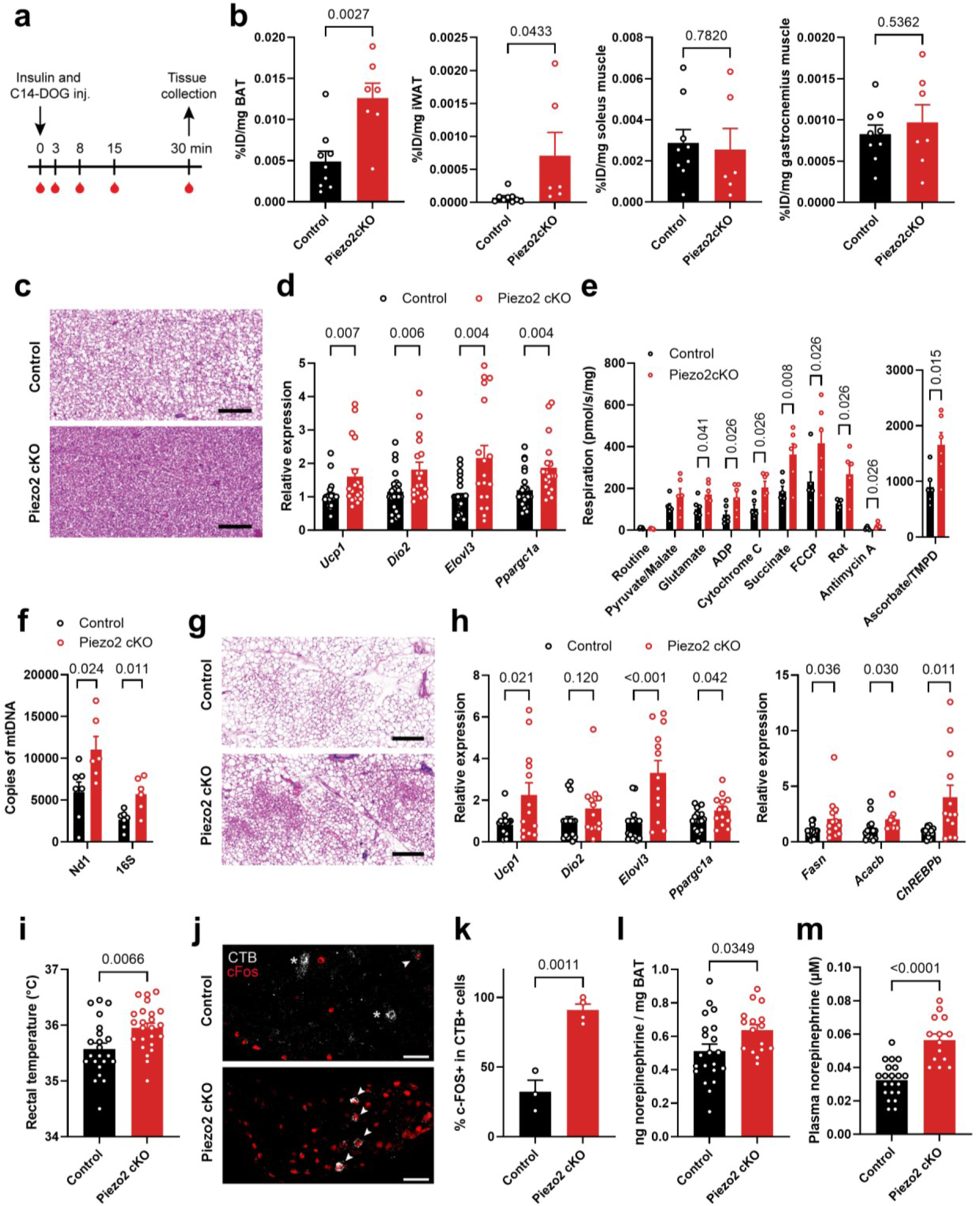
*Piezo2* cKO mice show increased glucose uptake and upregulated thermogenic program in iBAT and iWAT, and display elevated sympathetic tone. a) Schematic of the radiolabeled ITT experiment, in which insulin and C14 2-DG were injected together. Blood was collected from the tail at different time points. Mice were sacrificed 30 min after the injection and different tissues were collected. b) Elevated C14 2-DG uptake relative to littermate controls (n = 9) was observed in iBAT and iWAT of *Piezo2* cKO mice (n = 6-7), but not in soleus and gastrocnemius muscles. Data are presented as % of injected dose (ID) per mg tissue. c) H&E staining of mutant and control iBAT; scale bar 200 µm. d) Expression analysis of genes involved in thermogenic programs, including *Ucp1*, *Dio2, Elovl3* and *Ppargc1a,* in iBAT from *Piezo2* cKO mice (n = 18) normalized to controls (n = 23) using the 2^−ΔΔCt^ method. Statistical analysis was performed using multiple t-tests. e) Mitochondrial respiration was measured in iBAT using high-resolution respirometry. Detailed respiration measurements were made during basal respiration (routine), complex I-linked leak respiration (pyruvate, malate, glutamate), ADP stimulated (ADP), complex I maximal respiration (cytochrome C), OXPHOS capacity (succinate), maximal respiration (FCCP), complex I inhibition demonstrating complex II-linked respiration (rot), complex III inhibition demonstrating residual oxygen consumption (antimycin A) and complex IV respiration (Ascorbate and TMPD). n = 6 mice per group. Statistical analysis was performed using multiple Mann-Whitney tests. f) Increased copy number of mitochondrial DNA (mtDNA) in iBAT of *Piezo2* cKO mice compared to controls. Statistical analysis was performed using multiple t-tests. n=6 *Piezo2* cKO mice, n=7 littermate controls. g) H&E staining of iWAT; scale bar, 200 µm. h) Expression analysis of genes involved in thermogenic programs, including *Ucp1*, *Dio2, Elovl3* and *Ppargc1a,* and in de novo lipogenesis, including *Fasn*, *Acacb* and *ChREBPb,* in iWAT from *Piezo2* cKO mice (n = 13) normalized to controls (n = 16) using the 2^−ΔΔCt^ method. Statistical analysis was performed using multiple t-tests. i) Rectal temperature was measured in *Piezo2* cKO mice (n = 22) and controls (n = 24). j) Representative images of immunofluorescent staining for c-FOS in stellate ganglia from *Piezo2* cKO and control mice, with the retrograde tracer CTB injected into iBAT. Arrowheads indicate cells double-positive for c-FOS and CTB, while asterisks mark CTB-positive but c-FOS-negative cells. Scale bar, 50 µm. k) Quantification of c-FOS positive cells among CTB-labeled cells of the stellate ganglia in *Piezo2* cKO (n = 4 mice, 8 ganglia, 84 CTB-labeled cells) and control mice (n = 3 mice, 6 ganglia, 85 CTB-labeled cells). l) Norepinephrine levels in iBAT of *Piezo2* cKO mice (n=16) and littermate controls (n=21). m) Concentration of norepinephrine in plasma from *Piezo2* cKO mice (n = 14) and littermate controls (n = 21). Mice were kept at room temperature of around 22°C. Unless indicated otherwise, statistical analysis was performed using unpaired t-tests. Data are presented as mean ± SEM.

Typically, an increase in energy utilization in adipose tissue occurs during fat browning (*36*). Hence, we proceeded with histological and gene expression analyses of adipose tissues and found clear differences in H&E staining between iBAT from *Piezo2* cKO mice and controls (Fig. 5c). The expression of genes involved in the thermogenic program, including *Ucp1*, *Dio2*, *Elovl3* and *Ppargc1a*, was upregulated (Fig. 5d). Moreover, since adipose tissue function relies on mitochondria (*6*), we employed high-resolution respirometry (Oroboros O2k) to assess mitochondrial respiration in iBAT. We found a global increase in mitochondrial respiration in the iBAT of *Piezo2* cKO mice compared to controls. This included a significant increase in complex I-linked respiration (using pyruvate, malate, and glutamate (PMG)), adenosine diphosphate (ADP)-stimulated respiration, complex II-linked respiration (using succinate), FCCP-stimulated maximum respiration, and complex IV-linked respiration (using ascorbate and TMPD) (Fig. 5e). This increased mitochondrial respiration could be attributed to either a higher mitochondrial mass or an elevated mitochondrial activity. Hence, we assessed mitochondrial DNA (mtDNA) copy number by determining mtDNA to nuclear DNA ratio (*37*), and found that iBAT of *Piezo2* cKO mice contained a higher copy number of mtDNA relative to the controls (Fig. 5f). This indicates that the increased respiration is likely due to an increased number of mitochondria. Furthermore, we found a similar phenotype in iWAT of *Piezo2* cKO mice, that exhibited marked differences in H&E staining suggesting beiging and also showed reduced lipid droplet size (Fig. 5g and Fig. S8). Additionally, in mutant iWAT we observed an upregulated expression of *Ucp1*, *Elovl3* and *Ppargc1a* and of genes associated with de novo lipogenesis, including *Fasn*, *Acacb* and *ChREBPb* (Fig. 5h). An increase in lipogenesis during elevated thermogenic activity is a known effect that ensures fuel availability (*38*). In line with these findings, we found an elevated rectal temperature in *Piezo2* cKO mice (Fig. 5i). Collectively, these results suggest that Piezo2 in PV sensory neurons regulates morphological and physiological properties of iBAT and iWAT.

Adipose tissue browning/beiging is typically triggered by the sympathetic system through the release of norepinephrine (*12, 38*). To test the possibility that Piezo2 in PV sensory neurons modulates the sympathetic tone, we performed immunostaining for the neuronal activity marker c-FOS in stellate ganglia, whose sympathetic fibers innervate iBAT and other tissues (*39*). We marked sympathetic neurons innervating iBAT by injecting the retrograde tracer cholera toxin subunit B (CTB) in iBAT of *Piezo2* cKO mice and littermate controls. We found a significant increase in the number of c-FOS positive cells among the CTB-labeled population (Fig. 5j,k).

Furthermore, norepinephrine levels were increased in iBAT as well as in the plasma of *Piezo2* cKO mice (Fig. 5l,m). These findings support the possibility that Piezo2 in PV sensory neurons regulates norepinephrine secretion by the sympathetic nervous system, providing a potential mechanistic explanation for the observed adipose tissue and systemic phenotypes.

Given the role of the gastrointestinal system in whole-body metabolism (*40*) and the finding that Piezo2 controls gastrointestinal transit (*41*), we verified if deleting Piezo2 in PV sensory neurons affects gastrointestinal function. We found no overt sign of gastrointestinal dysfunction in *Piezo2* cKO mice (Fig. S9).

Together, our data reveal that the deletion of Piezo2 in PV sensory neurons leads to a phenotype in iBAT and iWAT, which exhibited increased glucose uptake while other tissues, including muscles, showed no increase. This suggests that brown and beige fat are likely driving the observed systemic metabolic phenotype.

### iBAT and iWAT are innervated by TrkC/Runx3/PV sensory neurons expressing Piezo2

Our findings suggest an important role for Runx3/PV sensory neurons and for Piezo2 in these neurons in regulating systemic and brown and beige fat metabolism. Adipose tissues are innervated by sensory neurons (*16*), yet their identity has remained largely elusive. Thus, we wondered if Runx3/PV sensory neurons, which are part of the TrkC subpopulation of somatosensory neurons (*20*), directly innervate these adipose tissues. To test this possibility, we used a retrograde labelling approach and injected cholera toxin subunit B (CTB) into iBAT and iWAT, and assessed the labelling in the dorsal root ganglia (DRG), where soma of sensory neurons reside (Fig. 6a). In rodents, sensory innervation in iBAT and iWAT arrives mainly from C4-T3 DRG and T11-L3 DRG, respectively (*16, 42*). We found that CTB signals largely co- localized with TrkC (Fig. 6b), In T2-T3 DRG, which innervate iBAT, we found that 84% of the CTB signals co-localized with TrkC (Fig. 6c). Similarly, in iWAT-innervating T12-L1 DRG, co- localization was 74% (Fig. 6c). We next tested whether adipose sensory neurons are also positive for PV and Runx3 by injecting CTB into iBAT and iWAT of *PV-Cre;tdTomato* mice and by assessing CTB, tdTomato and Runx3 labelling in DRG. We found that 42% and 35% of the CTB signals co-localized with both PV and Runx3 in T2-T3 DRG and T12-L1 DRG, respectively (Fig. 6d,e). We next verified this finding using a whole-body immunolabeling approach (*43*), which allows tracing axons and identifying their targets (*44*). Light-sheet imaging of cleared *PV-Cre;Brainbow* mouse (*44*) showed sensory neurons that originated in T2 and T3 DRG and projected into iBAT (Fig. 6f,g). Furthermore, stainings of iBAT sections confirmed that PV-positive neurons innervate the tissue (Fig. S10). Together, these data indicate that iBAT and iWAT are innervated by TrkC-, Runx3- and PV-positive sensory neurons.

**Fig. 6.**
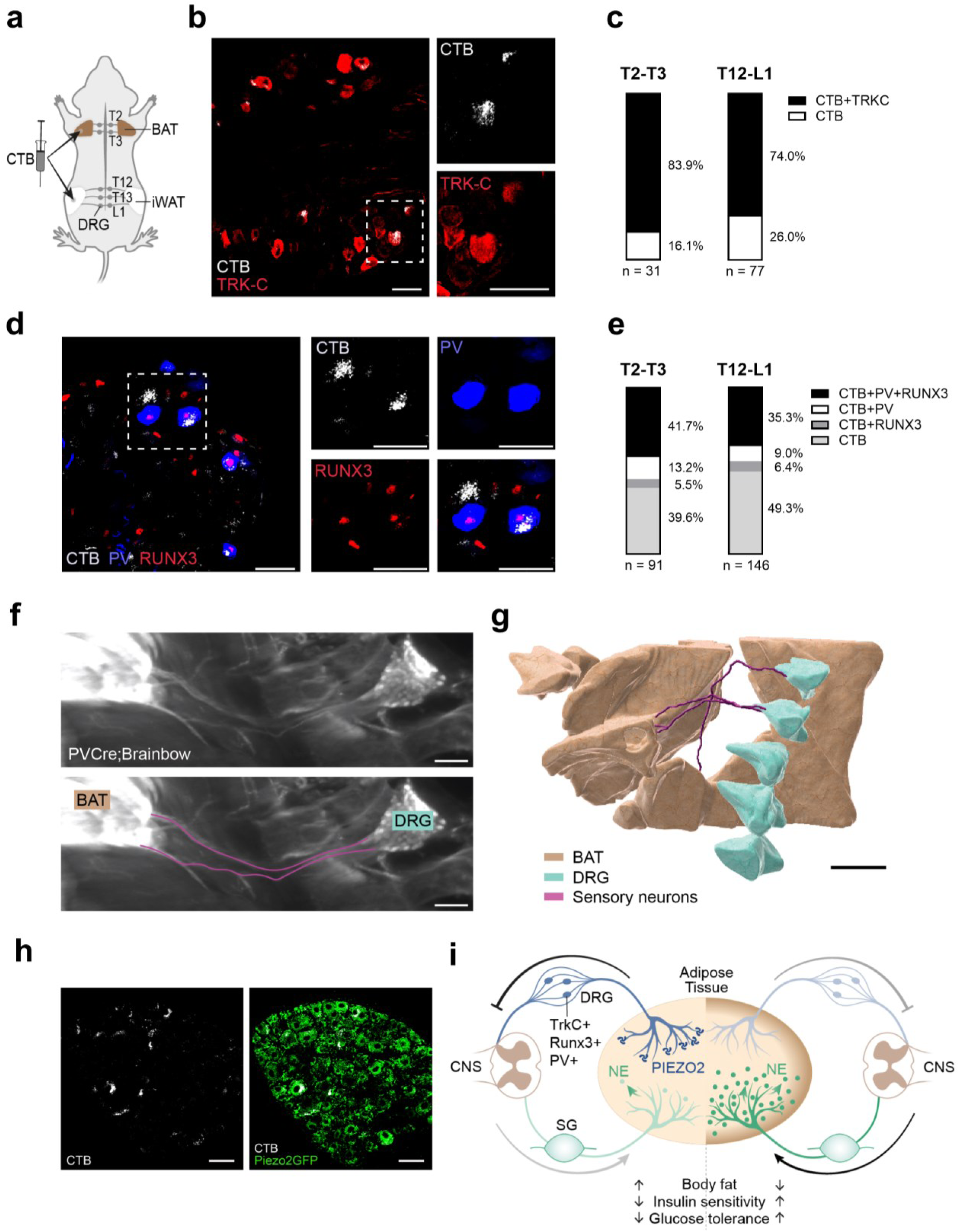
iBAT and iWAT are innervated by TrkC/Runx3/PV sensory neurons expressing Piezo2. a) Schematic of the retrograde tracing experiment, in which CTB was injected into iBAT and iWAT of a mouse. b) Representative image of a CTB-labeled T3 DRG section stained for TRKC. On the right are magnifications of the dashed square. Scale bars, 50 µm. c) Quantification of CTB- and TrkC-positive cells in DRG sections at indicated spine levels (n is the number of CTB signals from 3 mice). d) Representative image of a CTB-labeled T3 DRG section from a *PV-Cre;tdTomato* mouse stained for RUNX3. On the right are magnifications of the dashed square, with the bottom right image showing merger of the three signals. Scale bars, 50 µm. e) Quantification of CTB-, PV- and Runx3-positive cells in DRG sections from *PV- Cre;tdTomato* mice at indicated spine levels (n is the number of CTB signals from 3 mice). f) Images from whole-body immunolabeling of cleared *PV-Cre;Brainbow* mouse showing projections from T3 DRG to iBAT. Scale bar, 200 µm. g) 3D rendering of iBAT, DRG and sensory neurons from whole-body images of *PV-Cre;Brainbow* mouse. The top DRG is at the T2 level. Scale bar, 700 µm. h) Representative image of CTB labelling in a section of a T13 DRG, which innervates iWAT, in *Piezo2-GFP* mice. Scale bar, 50 µm. i) Proposed mechanism of systemic metabolism regulation by PIEZO2 in TrkC/Runx3/PV adipose sensory neurons.

In addition to the important role of Runx3/PV sensory neurons, we found that the mechanosensor Piezo2 is a key component in the mechanism that regulates adipose and systemic metabolism. To investigate if adipose sensory neurons express Piezo2, we injected CTB in iBAT and iWAT of *Piezo2-GFP* mice. Examination of DRG sections revealed co-localization of CTB and GFP signals (Fig. 6h and Fig. S11), confirming *Piezo2* expression in adipose sensory neurons. Of note, using single-molecule in situ hybridization chain reaction (HCR) (*45*), we found that *Piezo2* expression in iWAT sensory neurons is increased after 8 weeks of HFD (Fig. S12). Taken together, our study suggest that brown and beige fat are innervated by Runx3/PV sensory neurons, whose ablation leads to a systemic metabolic phenotype. Deletion of *Piezo2* in these neurons reproduces the phenotype and leads to elevated glucose uptake and *Ucp1* expression in brown and beige fat.

## Discussion

In this study, we uncovered that Runx3/PV sensory neurons are associated with the suppression of thermogenic programs in both brown and beige fat. Notably, we found that the expression of the mechanosensitive channel Piezo2 by sensory neurons is essential for this suppression, suggesting that the signal activating these neurons might be of a mechanical nature. Our discovery points to the suppression of norepinephrine secretion by the sympathetic system as the underlying mechanism. Lastly, we have established that this regulatory mechanism is key for controlling systemic metabolism in both physiological and pathological conditions. The concept of a sympathetic-sensory feedback loop between fat and brain is based on a series of experiments (*12, 14, 46*). For instance, sympathetic stimulation of adipose tissue induced a sensory response as observed by an increased neuronal activity in DRG (*14*). In addition, a viral tracing approach uncovered sympathetic-sensory circuits between brown fat and specific brain areas (*14*). Nonetheless, the function of this potential feedback loop remains poorly understood. Our study suggests the possibility that this feedback loop may play a regulatory role in preventing excessive fuel utilization, potentially acting as a sensory brake on the sympathetic system. In this context, our finding that temperature, which can activate the sympathetic system, also influences the sensory-mediated metabolic phenotype highlights the possibility that this feedback loop is temperature dependent. We found that a loss of sensory signaling, by ablating either sensory neurons or *Piezo2* expression, activates the sympathetic system, upregulates adipose thermogenic programs and causes significant systemic metabolic changes (Fig. 6i). Additional support for the role of sensory neurons as a negative feedback regulator comes from a recent study, in which local ablation of iWAT innervating sensory neurons resulted in sympathetic-mediated beiging (*16*). These findings raise several questions. For instance, the projection targets of these DRG neurons and the circuits they engage with remain unclear. Additionally, how the sensory information is integrated, processed, and translated into a functional output remains to be determined.

The discovery that sensory neurons negatively regulate the activation of brown and beige fat raises the intriguing question of the signal triggering the sensory response. Our identification of Piezo2 expression in sensory neurons innervating adipose tissue, coupled with the observation that blocking *Piezo2* expression leads to the activation of brown and beige fat, suggests that this molecule is a key mediator of the sensory signal. Given that Piezo2 is a mechanosensitive ion channel (*25, 26*), the signal activating sensory neurons is likely mechanical. Interestingly, exposure of mice to low temperatures was found to increase the stiffness of BAT relative to mice kept at room temperature (*47*). In line with this, β-adrenergic stimulation of brown adipocytes induces actomyosin-mediated contractile signaling, as observed in muscle tissues, which generates tension that is necessary for activating thermogenesis (*47*). These findings suggest that the biomechanical properties of adipose tissue are coupled to its metabolic state. We speculate that increases in local tissue tension or stiffness upon sympathetic-mediated fat activation might be detected by adipose sensory neurons via Piezo2, which might thereby convey the metabolic state of fat to the central nervous system. It will be of interest in future studies to test these hypotheses and investigate the cues activating adipose sensory endings and the role of Piezo2 in this process.

In metabolic disorders and aging, adipose tissues exhibit major changes, characterized by excessive lipid accumulation, insulin resistance, and pro-inflammatory, pro-fibrotic states with aberrant accumulation of extracellular matrix components, such as collagen VI, SPARC, thrombospondin-1 and hyaluronic acid (*48–52*). It will be important to investigate the role of adipose sensory neurons in metabolic disorders and aging, and how these conditions affect sensory function. We found an elevated Piezo2 expression in sensory neurons of obese iWAT, which may contribute to a vicious cycle, as increased sensory activity likely suppresses sympathetic-mediated adipose thermogenesis.

Thus far, understanding of the global effects of the sensory system in adipose tissue and in systemic metabolism has been limited. The molecular profiling of sensory neurons in adipose tissue in our study provides a molecular entry point to address these questions. However, this approach has limitations. One concern is that the markers we identified for adipose sensory neurons are shared with sensory neurons innervating the proprioceptors muscle spindles and Golgi tendon organs, which are located in muscles and tendons. This raises the immediate questions of whether altered proprioceptive sensory activity in our mouse models might be contributing to the phenotype and whether some of the effects observed involve other tissues, such as skeletal muscles. We addressed this with a comprehensive analysis of skeletal muscles using bulk RNA sequencing, fiber typing, and in vivo glucose uptake measurements. The results showed no overt differences in mutant muscles, supporting our conclusion that the main source for the observed systemic metabolic phenotype is the regulation of brown and beige fat by sensory neurons. This is also in line with the well-established role of adipose tissues in regulating systemic metabolism, including glucose homeostasis and insulin sensitivity (*6*). Nonetheless, identifying additional genetic approaches targeting adipose sensory neurons will be of interest.

In summary, this work identifies the sensory system and the mechanosensitive ion channel Piezo2 as central regulators that attenuate adipose glucose uptake and *Ucp1* expression and prevent systemic hypermetabolism. These findings offer a new direction for the development of therapeutic strategies for many who suffer from obesity, metabolic diseases as type 2 diabetes, and related comorbidities.

## Limitations of the study

Although we thoroughly characterized muscles of *Piezo2* cKO mice and found no evidence of a muscle involvement in the systemic metabolic phenotype, these mutants lack Piezo2 in proprioceptive sensory neurons innervating muscle spindles and Golgi tendon organs (*53*).

Hence, we cannot exclude a potential contribution of proprioceptive sensory neurons to the systemic metabolic phenotype, and further studies will be needed to delete Piezo2 in adipose sensory neurons while sparing proprioceptive ones.

## Acknowledgments

We thank the members of the Zelzer lab for insightful discussions. We thank Dr. Noa Wigoda from the Department of Life Sciences Core Facilities and Dr. Tsviya Olender from the Department of Molecular Genetics for bioinformatic analyses; Dr. Yoram Groner from the Department of Molecular Genetics for kindly providing Runx3 mice and antibodies for RUNX3 and TrkC; Dr. Ori Brenner from the Department of Veterinary Resources for pathological assessments, Yehuda Moshayev from the Safety Unit for assistance in handling radioactive material; Dr. Ron Rotkopf from the Department of Life Sciences Core Facilities for statistical assistance and Hanna Vega from the Design, Photography and Printing Branch for designing illustrations (all from the Weizmann Institute of Science). We thank Dr. Thomas Misgeld from the Institute of Neuronal Cell Biology, Technical University of Munich, for kindly providing Brainbow mice. We thank the de Picciotto Cancer Cell Observatory in memory of Wolfgang and Ruth Lesser, Weizmann Institute of Science, for providing light sheet imaging infrastructure. We thank Dr. Carla Horvath from the Institute of Food, Nutrition and Health at ETH Zurich for insightful discussions and advices. We thank Nitzan Konstantin for editorial assistance. F.S.P was supported by a SNSF Postdoc Mobility fellowship, a fellowship from the Swiss Friends of the Weizmann Institute of Science and an EMBO non-stipendiary Postdoctoral fellowship. K.D.B. is endowed by the Schulthess Foundation. Y.K. is the Incumbent of the Sarah and Rolando Uziel Research Associate Chair.

## Authors contributions

F.S.P conceptualized the study, designed and performed the experiments and wrote the manuscript. B.B. performed staining experiments and contributed to data interpretation, experimental design and drafting the manuscript. S.R. performed staining experiments, light-sheet imaging, and segmentation analyses. Y.K. performed and supervised metabolic experiments, and contributed to data interpretation. S.K. performed radiolabeled glucose uptake and in situ hybridization experiments. E.M. performed muscle staining experiments. T.M. and A.B. performed norepinephrine measurements. Y.A. performed and supervised light-sheet imaging experiments. S.H-O.S. and T.Y. performed and supervised mitochondrial respiration experiments. E.A.R., A.T. and K.D.B. supervised metabolic analyses and contributed to experimental design, data analysis and interpretation. E.Z. supervised the study, designed experiments, acquired funding, and wrote the manuscript. All authors approved the final version of the paper.

## Resource availability

### Lead contact

Resources or reagents should be requested from the lead contact, Elazar Zelzer.

### Materials availability

This study did not generate new unique reagents.

## Data and code availability

- Bulk RNA-seq data can be found at the Gene Expression Omnibus under the accession number GSE256369.
- Data presented in this manuscript are provided in Data S1.
- This paper does not report original code.
- All other data are available from the corresponding author on request.

## Star Methods

### Key resource table

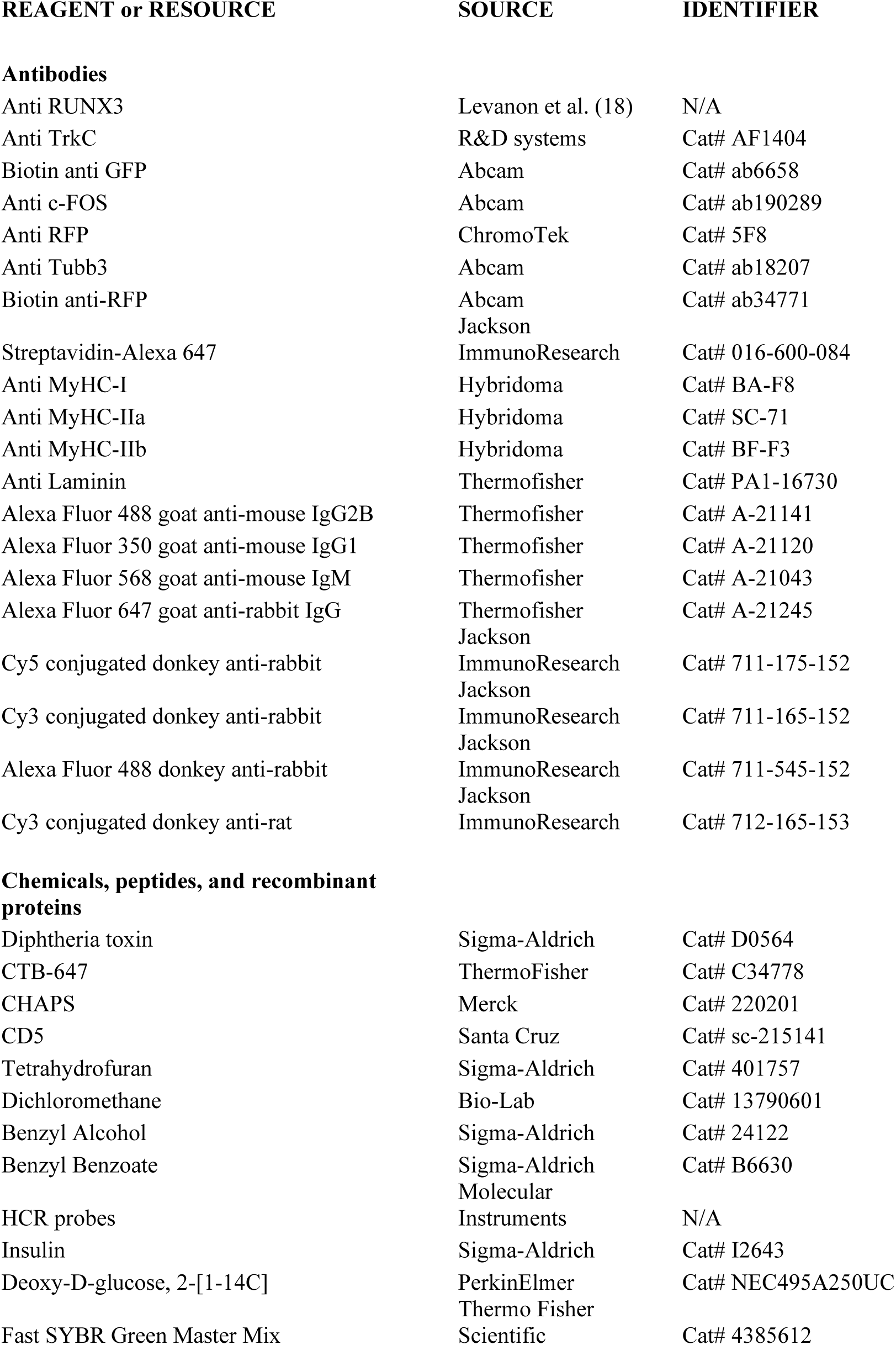
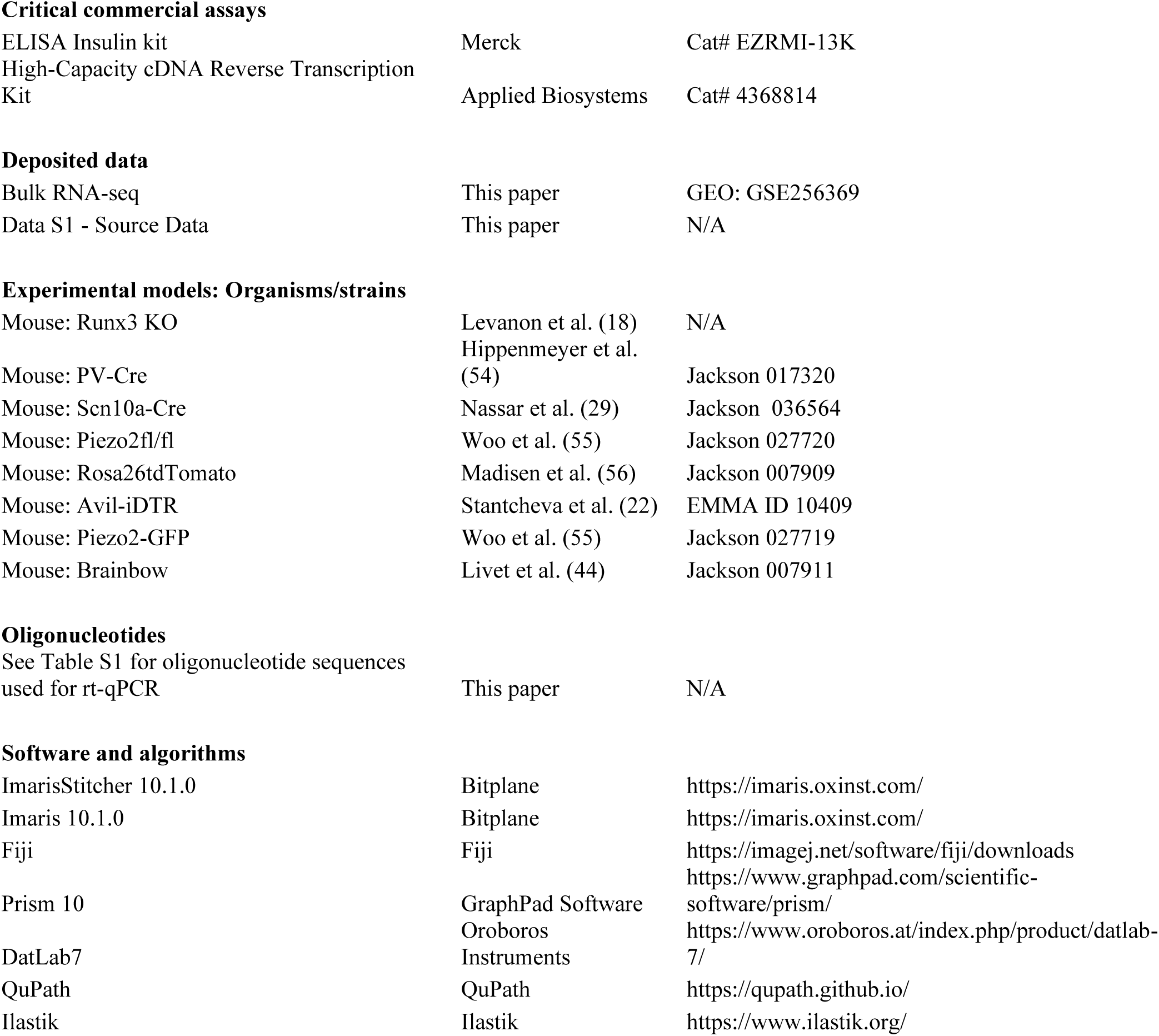

### Experimental model and study participant details Mice

The generation of *Runx3* KO (*Runx3*-null, ICR background) (*18*), *PV-Cre* (*parvalbumin-Cre*, C57BL/6J background; Jackson 017320) (*54*), *Scn10a-Cre* (C57BL/6 background, Jackson 036564) (*29*), *Piezo2^fl/fl^* (loxP-flanked (floxed) *Piezo2*, C57BL/6J background; Jackson 027720) (*55*), *Rosa26^tdTomato^* (*Gt(ROSA)26Sor^tm9(CAG-tdTomato)Hze^*, C57BL/6J background; Jackson 007909) (*56*), Avil-iDTR (*advillin^iDTR^*, C57BL/6 background; EMMA ID 10409)(*22*), *Piezo2-GFP* (*Piezo2-GFP-IRES-Cre*, C57BL/6J background; Jackson 027719) (*55*), and Brainbow mice (*Thy1-Brainbow1.1*, C57BL/6 background, Jackson 007911; kindly provided by Dr. Thomas Misgeld from the Technische Universität, München) (*44*) was described previously. *Runx3* KO mice were generated by crossing *Runx3* heterozygous males and females. Littermates that were either wild type or *Runx3^+/^*^-^ were used as controls. *PV-Cre;Piezo2^fl/fl^* mice were generated by crossing males carrying *PV-Cre* and a single *Piezo2*-floxed allele (LoxP/+) with females homozygous for the *Piezo2* loxP allele; *Piezo2^fl/fl^*and *PV-Cre;Piezo2^fl/+^* littermates were used as controls. *Scn10a-Cre;Piezo2^fl/fl^* mice were generated by crossing males heterozygous for *Scn10a- Cre* and heterozygous for the *Piezo2* loxP allele with females homozygous for the *Piezo2* loxP allele; *Piezo2^fl/fl^* littermates were used as controls. *PV-Cre;Avil-iDTR* mice were generated by crossing males carrying *PV-Cre* with females heterozygous for *Avil-iDTR*. Littermates carrying either *Adv-iDTR* or *PV-Cre* were used as controls. *PV-Cre;Rosa26^tdTomato^* and *PV-Cre;Brainbow* reporter mice were generated by crossing males carrying *PV-Cre* with females homozygous for *Rosa26^tdTomato^* and heterozygous for Brainbow, respectively.

Unless indicated otherwise, all experiments were performed with adult female and male mice of the following ages: 2-5 months old *Runx3 KO* mice; 3-6 months old *PV-Cre;Piezo2^fl/fl^* mice; 3-6 months old *Scn10a-Cre;Piezo2^fl/fl^* mice; *PV-Cre;Avil-iDTR* mice were 2-3 months old at the time of DT injection and metabolic tests were performed 1-2 months later. Mice were kept in standard housing at room temperature (around 22°C) with a 12/12 h light-dark cycle. Mice were fed ad libitum with a chow diet or a 60% high-fat diet. All experiments were approved by the Institutional Animal Care and Use Committee (IACUC) of the Weizmann Institute.

## Method details

### Diphtheria toxin administration

Diphtheria toxin (D0564, Sigma-Aldrich) was injected intraperitoneally at a concentration of 150 µg/kg body weight in 0.9% saline.

### Retrograde labelling

Mice were anesthetized with ketamine/xylazine (100 mg/kg body weight and 20 mg/kg body weight, respectively) and given ophthalmic ointment and buprenorphine (0.05 mg/kg body weight). The skin was shaved and sterilized at the surgical areas. Then, an incision in the skin was made at the interscapular region for injections into the iBAT. For iWAT injections, incisions in the skin were made on the flank of both sides. Based on a previous report (*16*), for retrograde labelling we injected CTB-647 (ThermoFisher, C34778) at either 0.1 µg/µl or 0.01 µg/µl in PBS using a 5 µl Hamilton syringe. A volume of 2.5 µl was injected into each side of the iBAT and a volume of 5 µl was injected into each iWAT, both through multiple injections. After the injections, the skin was sutured. Mice were euthanized 4-5 days after the injection and stellate ganglia or DRG were isolated freshly.

### Immunostaining of cryosections

DRG were isolated as described previously (*57*), fixed overnight in 4% paraformaldehyde (PFA)/PBS at 4°C, transferred to 30% sucrose overnight, and then embedded in OCT and sectioned by cryostat at a thickness of 20 µm. Next, cryosections were dried for 30 min at room temperature, permeabilized with 0.3% Triton X-100 in PBS, washed with 0.1% Triton X-100 in PBS for 5 min and blocked in 5% horse serum, 1% bovine serum albumin (BSA), and 0.05% Triton X-100 in PBS for 1 h. Then, sections were incubated overnight at 4°C with the following primary antibodies: anti-RUNX3 (rabbit, 1:800, (*18*)), anti-TrkC (goat, 1:250, (*58*)) or anti-GFP (Biotin goat, Abcam ab6658, 1:200). The next day, sections were washed three times in 0.1% Triton X-100 in PBS and incubated for 1 h with secondary fluorescent-conjugated antibody, washed three times in 0.1% Triton X-100 in PBS, counterstained with DAPI and mounted with Immu-mount mounting medium (Thermo Fisher Scientific).

For stellate ganglia immunofluorescence, freshly dissected ganglia (*59*) were immediately embedded in OCT and sectioned by cryostat at a thickness of 20-40 µm. c-FOS staining was performed as described previously (*60*). In brief, sections were fixed in 2% PFA/PBS for 5 min, washed in PBS with 0.3% Triton X-100, and incubated with 0.1 M glycine for 5 min, washed and permeabilized with PBS with 0.3% Triton X-100 for 20 min. Slides were then blocked with 5% donkey serum, 1% BSA, and 0.3% Triton X-100 in PBS for 2 h at room temperature and incubated with primary c-FOS antibody (rabbit, Abcam ab190289, 1:1000) overnight at 4°C. Then, slides were washed with PBS with 0.3% Triton X-100, incubated with secondary antibody for 2 h at room temperature and washed four times with PBS with 0.3% Triton X-100, counterstained with DAPI and mounted with Immu-mount.

For adipose tissue immunostaining, adult *PVCre;Rosa26^tdTomato^*mice were deeply anesthetized and perfused intracardially with cold, heparinized PBS (10 U/ml heparin, Sigma H3149) followed by cold 4% PFA in heparinized PBS using a peristaltic pump. The skin was removed and the mouse body postfixed in 4% PFA in PBS for 6 h at 4°C, followed by washes in PBS. iBAT was dissected and transferred to 30% sucrose overnight, and then embedded in OCT and sectioned by cryostat at a thickness of 30 µm. Next, cryosections were dried for 90 min at room temperature, fixed in 4% PFA/PBS for 10 min, permeabilized with 0.3% Triton X-100 in PBS, washed with 0.1% Triton X-100 in PBS for 5 min, bleached with 10% H_2_O_2_ in Methanol for 2 h, and blocked with CAS-Block™ (Thermo Fisher Scientific). Slides were then incubated with primary antibodies anti-RFP (rat, 1:100, ChromoTek [5F8]) and anti-Tubb3 (rabbit, 1:500 Abcam ab18207) overnight at 4°C. Then, slides were washed with PBS with 0.1% Triton X-100, incubated for 1 h with secondary fluorescent-conjugated antibody, washed three times in 0.1% Triton X-100 in PBS, counterstained with DAPI and mounted with Immu-mount mounting medium (Thermo Fisher Scientific).

Sections were Imaged using Zeiss LSM800 or LSM900 confocal microscope. Images were processed with Fiji (*61*).

### Whole-body immunolabeling using wildDISCO

Whole-body immunolabeling of *PV-Cre;Brainbow* mice was performed using wildDISCO as described previously (*43*). In brief, P15 mice were deeply anesthetized and perfused intracardially with cold, heparinized PBS (10 U/ml heparin, Sigma H3149) followed by cold 4% PFA in heparinized PBS using a peristaltic pump. The skin was removed and the mouse body postfixed in 4% PFA in PBS for 6 h at 4°C, followed by washes in PBS. Subsequently, the tip of the perfusion needle was inserted into the heart through the hole made in the initial perfusion and glued to ensure stable perfusion. Mouse body was perfused overnight with PBS, and then for 2 days with the decalcification solution (10% w/v EDTA in 0.01 M PBS; pH was adjusted to 8–9 with sodium hydroxide), followed by three washes of 3 h each with PBS. Then, the mouse body was perfused for 1 day with permeabilization and blocking solution (10% horse serum and 2% Triton X-100 in PBS) in 100 ml volume at room temperature, followed by 7 days treatment with primary antibodies diluted 1:10,000 in 100 ml immunostaining buffer (3% horse serum, 10% CHAPS (Merck, 220201), 2% Triton X-100, 10% DMSO, 1% glycine (MP, 808822), 1% CD5 (Santa Cruz, sc-215141) in PBS) at room temperature. Primary antibodies were biotin anti-RFP (Abcam, ab34771) and biotin anti-GFP (Abcam, ab6658). Then, 3 times 12 h of washes were carried out with PBS, followed by 7 days treatment with secondary antibody Streptavidin-Alexa 647 (Jackson ImmunoResearch, 016-600-084) diluted 1:10,000 in 100 ml of the same immunostaining buffer. Then, three 12-h washes with PBS were performed. Mice were then transferred to a chemical hood, placed on a gentle shaker and cleared using the following 3Disco passive whole-body clearing protocol: gradient of tetrahydrofuran (Sigma, 401757) in distilled water (50%, 70%, 80%, 100% twice), each step for 12 h with 60 ml volume, then 3 h in 60 ml dichloromethane (Bio-Lab, 13790601). Lastly, mice were kept in BABB (Benzyl Alcohol:Benzyl Benzoate at a ratio of 1:2, Sigma, 24122 and B6630) until the bodies were optically transparent. Light sheet fluorescent microscopy (Ultramiscroscope II, LaVision Biotec) was then performed using the ImspectorPro software (LaVision BioTec). The light sheet was generated by a Superk Super-continuum white light laser (emission 460 nm-800 nm, 1 mW/nm (NKT photonics), excitation filters 560/540 nm and 640/630 nm, emission filters 630/675 and 690/650). The microscope was equipped with a zoom body 2X objective and a corrected objective cap. Samples were glued to the sample holder and placed in an imaging chamber made of 100% quartz (LaVision BioTec) filled with BABB and illuminated from both sides by the laser. Images were acquired by an Andor Neo sCMOS camera (2,560 x 2,160, pixel size 6.5 μm x 6.5 μm, Andor – Oxford instruments) resulting in a pixel size of 3.0 μm XY. Z stacks were acquired with 4 μm steps and larger fields of view were imaged by tiling with 10% overlap. Images were stitched in ImarisStitcher 10.1.0 (Bitplane). Manual segmentation of DRG and iBAT from whole-body images were performed in Microscopy Image Browser (version 2.81) (*62*) and visualized in 3D in Imaris 10.1.0 (Bitplane). Segmentation of individual neurons was performed manually with simple neurite tracer in Fiji (*61, 63*).

### Single-molecule *in situ* HCR

DRG were isolated, fixed overnight in 4% paraformaldehyde (PFA)/PBS (DEPC) at 4°C, transferred to 30% sucrose overnight, and then embedded in OCT and sectioned by cryostat at a thickness of 10 µm. *Piezo2* HCR probe was designed for exons 43 to 45 and ordered from http://www.molecularinstruments.com. Cryosections were then processed for HCR smFISH following the manufacturer’s instructions. Briefly, slides were incubated with *Piezo2* probe overnight, washed and incubated with fluorophore-conjugated amplifiers. The sections were counterstained with DAPI, mounted with Immu-mount, and imaged using a LSM 800 confocal microscope (Zeiss). CTB-positive single cells were segmented and their *Piezo2* average signal intensity was measured in Fiji.

### Body composition analysis

Measurements of body fat and lean tissue were made in a sampling compartment using the Minispec LF50 Body Composition Analyzer (Bruker, USA).

### Metabolic cage studies

The PhenoMaster system (TSE-Systems, Germany), which contains sensitive feeding sensors and photobeam-based activity sensors, was used to assess oxygen consumption rate, carbon dioxide release rate, energy expenditure, food and water intake, as well as spontaneous locomotor activity. These data were acquired continuously and simultaneously in all cages. Mice were singly housed at room temperature of around 22°C and were allowed to habituate for at least two days before data acquisition. As previously suggested (*64*), data on oxygen consumption rate, carbon dioxide release rate and energy expenditure were analyzed using ANCOVA with body weight as a covariate.

### Glucose and insulin analyses

For glucose and insulin analyses, mice were kept at room temperature of around 22°C. To determine fasting blood glucose levels, glucose was measured in tail blood after 6 h of fasting. For each mouse, two measurements were made on different days and the results were averaged. For glucose tolerance test, mice were fasted for 6 h. Tail blood was collected at 0, 15, 30, 60, 90 and 120 min after an intraperitoneal injection of glucose (0113, J.T.Baker) at 2 or 1 g/kg body weight for chow-fed or HFD-fed mice, respectively.

For fasting plasma insulin, blood was collected from the tail using EDTA-coated capillaries (Microvette 200 K3E, Sarstedt) after 6 h of fasting. Blood was then centrifuged at 3000 g for 5 min and plasma was snap-frozen and stored at -80°C. Plasma insulin levels were analyzed using an ELISA Insulin kit (EZRMI-13K, Merck) following the manufacturer’s instructions.

For insulin tolerance test, mice were fasted for 3 h (chow diet) or 6 h (HFD). Tail blood was collected at 0, 15, 30, 60, 90 and 120 min after an intraperitoneal injection of insulin (I2643, Sigma-Aldrich) at 0.6 or 1 U/kg body weight for chow-fed or HFD-fed mice, respectively. In all tests, blood glucose was measured using testing strips and a blood glucometer (FreeStyle Optium Neo, Abbott).

### Housing at thermoneutrality

Starting shortly after birth (p1-p3), mice were moved into a temperature-controlled chamber (Tecniplast) with housing temperature of 30°C, corresponding to murine thermoneutrality (*31*).

### Histology

For hematoxylin and eosin (H&E) staining, tissues were fixed overnight in 4% PFA-PBS, dehydrated to 70% ethanol, embedded in paraffin and sectioned. Staining was carried out with a standard protocol.

### High-fat diet

Adult mice were fed with a high-fat diet (D12492i rodent diet with 60 kcal% fat, Research Diets, USA) for 12 weeks. Body weight and body composition were measured every 4 weeks. Glucose and insulin tolerance tests were carried out as described above during week 9-10 and week 10- 11, respectively. The bedding used was composed of a shredded natural hardwood (7093 Teklad Shredded Aspen Bedding).

### Tracing of radiolabeled 2-deoxyglucose uptake

Glucose uptake in various tissues was assessed by tracing radiolabeled 2-deoxyglucose (2-DG) as described previously (*65*). Mice were fasted for 4 h in single cages and subsequently injected intraperitoneally with deoxy-D-glucose, 2-[1-14C] (0.17 uCi/g body weight, PerkinElmer, NEC495A250UC) together with insulin (I2643, Sigma-Aldrich) at 1 U/kg lean body mass. Tail blood was collected at 0, 3, 8, 15 and 30 min for glucose and radioactivity measurements. 30 min after the injection, mice were euthanized by cervical dislocation and the following tissues were collected: iBAT, iWAT, eWAT, soleus, gastrocnemius, tibialis anterior, and extensor digitorum longus muscles, liver, heart, and brain. Radiolabeled 2-DG was measured in 20 µl of blood using scintillation counting. To assess the accumulation of phosphorylated radiolabeled 2-DG in each tissue, we used the precipitation method (*66*), followed by scintillation counting.

Tissues were isolated, snap-frozen in liquid nitrogen, stored at –80°C and subsequently homogenized with Trizol reagent (15596018, Invitrogen). RNA was isolated using MaXtract high-density tubes (129056, Qiagen) following the manufacturer’s protocol. The quality and quantity of the RNA were measured using a NanoDrop One spectrophotometer (Thermo Fisher Scientific). RNA was transcribed to cDNA using the High-Capacity cDNA Reverse Transcription Kit according to the manufacturer’s instructions (4368814, Applied Biosystems). Gene expression was analyzed by RT–qPCR with cDNA corresponding to 10 ng of starting RNA using the Fast SYBR Green Master Mix (4385612, Thermo Fisher Scientific). Samples were amplified with a StepOnePlus Real-Time PCR System (Applied Biosystems) using the following conditions: 10 min at 95 °C followed by 40 PCR cycles at 95 °C for 15 s and at 60 °C for 1 min. Relative gene expression levels were quantified using the 2^−ΔΔct^ method. *Tbp* and *Hprt1* were used as reference genes. A list of the primers is provided in Supplementary Table 1.

### Mitochondrial respiration measurement

Mitochondrial respiration was assessed using high resolution FluoRespirometry (Oroboros Instruments). Male mice were sacrificed using cervical dislocation and iBAT samples were weighted and immediately dissected and homogenized in MiR05 using Dounce homogenizers. Then the homogenized sample was added to the respirometry chambers. A substrate-uncoupler-inhibitor titration (SUIT-) protocol was used to evaluate mitochondrial respiration at different mitochondrial respiratory states. Titration protocol: pyruvate, malate, glutamate (5 mM, 2 mM, 10 mM), ADP (2.5 mM), cytochrome c (10 µM) succinate (10 mM), FCCP (repeated up to 4µM total maximal respiration), rotenone (0.5 µM), antimycin A (2.5 µM), ascorbate (2 mM), TMPD (0.5mM). After each addition, a plateau in respiration was awaited. Respirometry chambers were reoxygenated by opening for 5 min before the addition of FCCP or after ascorbate and TMPD. Data was analyzed using DatLab7 and normalized to the weight.

### Assessment of mitochondrial DNA copy number

Copy number of mitochondrial DNA in iBAT was assessed as previously described (*37*). Briefly, DNA was isolated using the DNeasy kit (69504, Qiagen) following the manufacturer’s protocol. RT–qPCR analysis was performed using 20 ng starting DNA and Fast SYBR Green Master Mix (4385612, Thermo Fisher Scientific) for the *Nd1* and *16S* genes belonging to mtDNA and for the nuclear encoded gene *Hk2*, which was used as reference. mtDNA copy number was calculated as 2x2^Δct^. A list of the primers is provided in Supplementary Table 1.

### Segmentation of lipid droplets in iWAT

To assess lipid droplet area, regions of interest from Panoramic slide scanner H&E images were selected in Qupath (*67*) and exported as tif files. Blood vessels were masked out using Ilastik (*68*) and images were segmented in Cellpose3 (*69*), using the cyto3 algorithm. Incorrect segmentations were manually removed from images and resegmented manually in Cellpose. Using Morpholib fiji plugin (*70*), adipocytes bordering the image edges were removed and lipid droplet area was quantified. iWAT from male mice were analyzed.

### Body temperature measurement

Core body temperature was measured using the rodent thermometer BIO-TK8851 (BiosebLab) with a thermocouple rectal probe. For each mouse, two measurements were taken in the middle of the light cycle on two different days and averaged.

### Norepinephrine measurements

Plasma norepinephrine level was measured as follows: First blood was collected retro-orbitally in the middle of the light cycle into MiniCollect tubes containing K3EDTA (450530, Greiner). Blood was centrifuged at 3000 g for 5 min and plasma was snap-frozen and stored at -80°C.

Next, a plasma aliquot of 10 µl was mixed with 40 µl methanol, vortexed and centrifuged to precipitate proteins (21,000 g for 10 min). A portion of the obtained supernatant (10 µl) was mixed with D6-norepinephrine (10 µl of 100 ng/ml) as internal standard and derivatized using Aqc reagent in 150 mM borate buffer. Tissue norepinephrine level was measured as follows: Adipose tissue was homogenized with acetonitrile (90 µl) and D6-norepinephrine (10 µl of 4 µg/ml) as internal standard using pestle motor grinder (Argos A0001), then incubated during 15 min (2,000 rpm, 10°C; Thermomixer C, Eppendorf), and centrifuged to precipitate proteins (21,000 *g*, 10 min). A portion of the obtained supernatant (10 µl) was derivatized using Aqc reagent in 150 mM borate buffer. The derivatized samples were analyzed by liquid chromatography-tandem mass spectrometry as described previously (*71*).

### Bulk RNA-seq of muscles

For sequencing, soleus and gastrocnemius muscles were isolated from male mice, snap-frozen and stored at -80°C. Subsequently, samples were homogenized with Trizol reagent (15596018, Invitrogen) and RNA was isolated using MaXtract high density tubes (129056, Qiagen) following the manufacturer’s protocol. RNA quality and quantity were assessed with a NanoDrop (Thermo Fisher Scientific) and a TapeStation (Agilent). RNA-seq libraries were prepared at the Crown Genomics institute of the Nancy and Stephen Grand Israel National Center for Personalized Medicine (INCPM), Weizmann Institute of Science. Libraries were prepared using the INCPM-mRNA-seq protocol. Briefly, the polyA fraction (mRNA) was purified from 500 ng of total input RNA followed by fragmentation and generation of double-stranded cDNA. After Agencourt Ampure XP beads cleanup (Beckman Coulter), end repair, A base addition, adapter ligation, and PCR amplification steps were performed. Libraries were quantified by Qubit (Thermo fisher scientific) and TapeStation (Agilent). Sequencing was done on a Nextseq instrument (Illumina) using a HO 75 cycles kit, allocating approximately 18M reads per sample (single-read sequencing). A user-friendly Transcriptome Analysis Pipeline (UTAP) version 1.10.2 was used for analysis (*72*). Reads were mapped to the *M. musculus* genome (UCSC, mm10) using STAR (v2.5.2b) (*73*) and GENECODE annotation. Only reads with unique mapping were further analyzed. Gene expression was calculated and normalized using DESeq2 version 1.16.1 (*74*), using only genes with a minimum of five reads in at least one sample. Raw p-values were adjusted for multiple testing (*75*). A gene was considered differentially expressed if it passed the following thresholds: minimum mean normalized expression of 5, adjusted p-value ≤0.05, and absolute value of log_2_ fold change ≥1.

### Muscle fiber typing

Fiber type staining was performed as previously described (*76*). In brief, soleus muscles were freshly isolated from male mice, embedded in Tissue-Tek and frozen in liquid N2-cooled isopentane. Frozen muscle cross sections (7-10 μm thick) were collected on Superfrost Ultra Plus slides (Thermo Fischer Scientific) using a cryostat (Leica CM 1950). Cross sections were permeabilized for 10 min in PBS with 0.5% Triton X-100, washed in PBS and blocked for 1 h at room temperature in PBS with 10% goat serum (16210064, Thermo Fisher Scientific).

Then, a primary antibody cocktail diluted in PBS with 10% goat serum was applied for 2 h against MyHC-I (1:50 dilution, BA-F8 from hybridoma, Iowa City, IA, USA), MyHC-IIa (1:200 dilution, SC-71 from hybridoma), MyHC-IIb (1:100 dilution, BF-F3 from hybridoma) and laminin (1:200 dilution, PA1-16730, Thermofisher). After washing three times for 5 min, a secondary antibody cocktail diluted in PBS with 10% goat serum was applied for 60 min.

Secondary antibodies included Alexa Fluor 488 goat anti-mouse IgG2B (1:250 dilution, A- 21141, Thermo Fisher Scientific), Alexa Fluor 350 goat anti-mouse IgG1 (1:250 dilution, A-21120, Thermo Fisher Scientific), Alexa Fluor 568 goat anti-mouse IgM (1:250 dilution, A- 21043, Thermo Fisher Scientific) and Alexa Fluor 647 goat anti-rabbit IgG (1:250 dilution, A- 21245, Thermo Fisher Scientific). After three washes of 5 min with PBS, slides were mounted and sealed with glass cover slips. Images of the muscle sections were captured using an epifluorescent microscope (Zeiss Axio observer Z.1, Zeiss, Germany) with a 10X objective, and stitched together using the tiles module in the ZEN 2011 imaging software (Zeiss). All images were taken with the same exposure time. Fiber diameter (minimum Feret diameter), fiber area and fiber type distribution were assessed by analyzing the images with Fiji (*61*) as described previously (*76, 77*).

### Stool analysis

Stool analysis was performed as previously described (*41*). In brief, stools were collected in the middle of the light cycle for 1 hour. Fresh stools were placed individually in Eppendorf tubes and weighed. Then, samples were dried overnight at 100°C and weighed again to determine dry weight and to calculate water content.

---

**Fig. S1.**
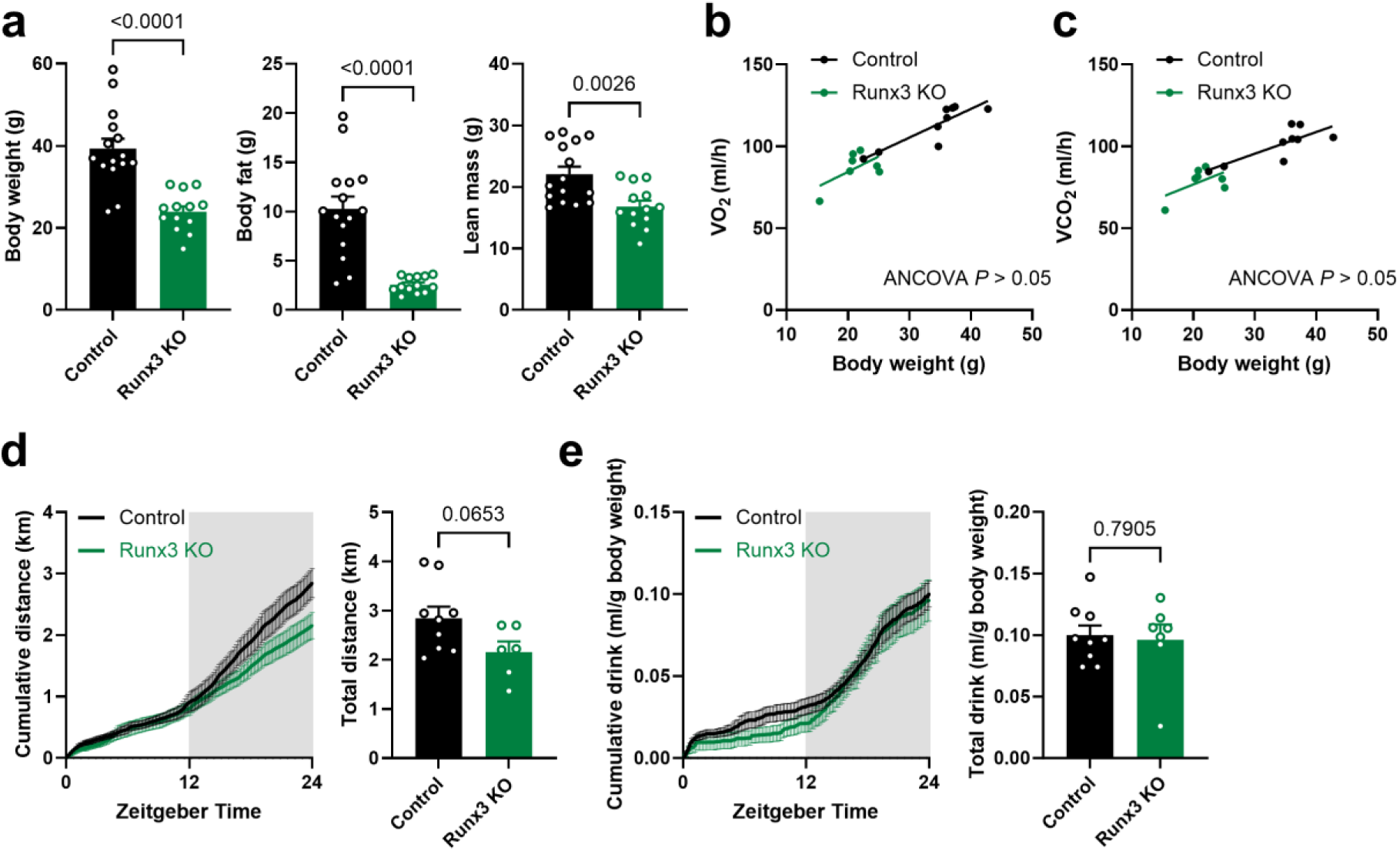
Body composition, calorimetric and behavioral assessment of *Runx3* KO mice. a) Body weight, body fat and lean mass of *Runx3* KO mice (n = 13) and littermate controls (n = 15). b-e) Metabolic cage analysis provided data on b) oxygen consumption (VO_2_), c) carbon dioxide release (VCO_2_), d) mobility and e) water intake. Mice were allowed to habituate in the metabolic cages for three days, after that data were acquired. Data are the average of two consecutive days. Mice were singly housed at room temperature of around 22°C. n=6-7 *Runx3* KO mice, n=9 littermate controls. Data in a, d and e were analyzed using an unpaired t-tests. Data in b,c were analyzed using ANCOVA with body weight as covariate. Data are mean ± SEM.

**Fig. S2.**
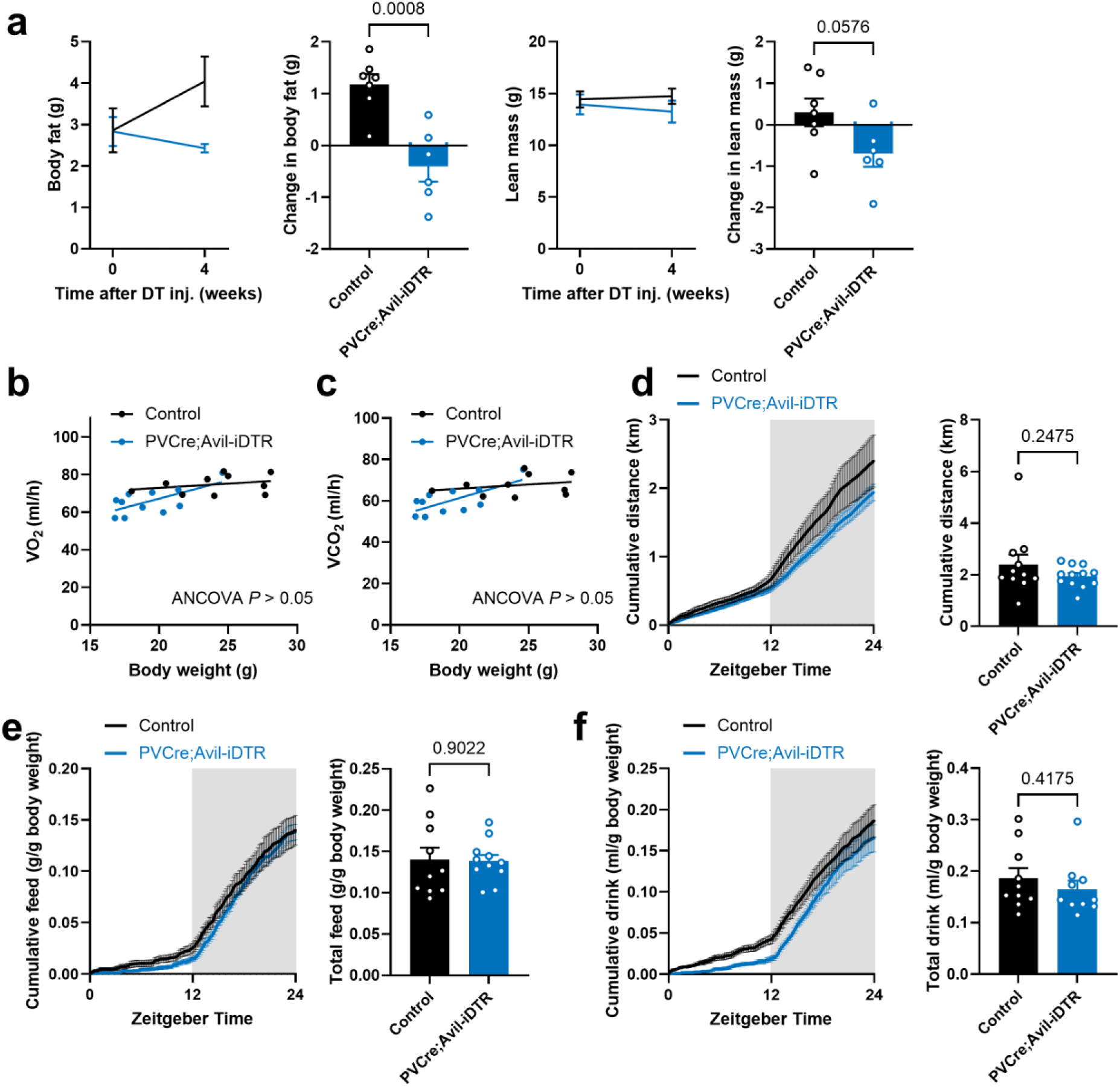
Body composition, calorimetric and behavioral assessment of *PVCre;Avil-iDTR* mice. a) Diphtheria toxin was injected into 2-3 months old *PV-Cre;Avil-iDTR* mice (n = 6) and control littermates (n = 7). Body composition was analyzed before and 4 weeks after the injection. b-f) Metabolic cage analysis was performed 4 to 8 weeks after the diphtheria toxin injection and provided data on b) oxygen consumption (VO_2_), c) carbon dioxide release (VCO_2_), d) locomotor activity, e) food and f) water intake. Mice were allowed to habituate in the metabolic cages for two days, after that data were acquired. n=11 *PVCre;Avil-iDTR* mice, n=10 littermate controls. Data are the average of two consecutive days. Mice were singly housed at room temperature of around 22°C. Data in a and d-f were analyzed using an unpaired t-tests. Data in b,c were analyzed using ANCOVA with body weight as covariate. Data are mean ± SEM.

**Fig. S3.**
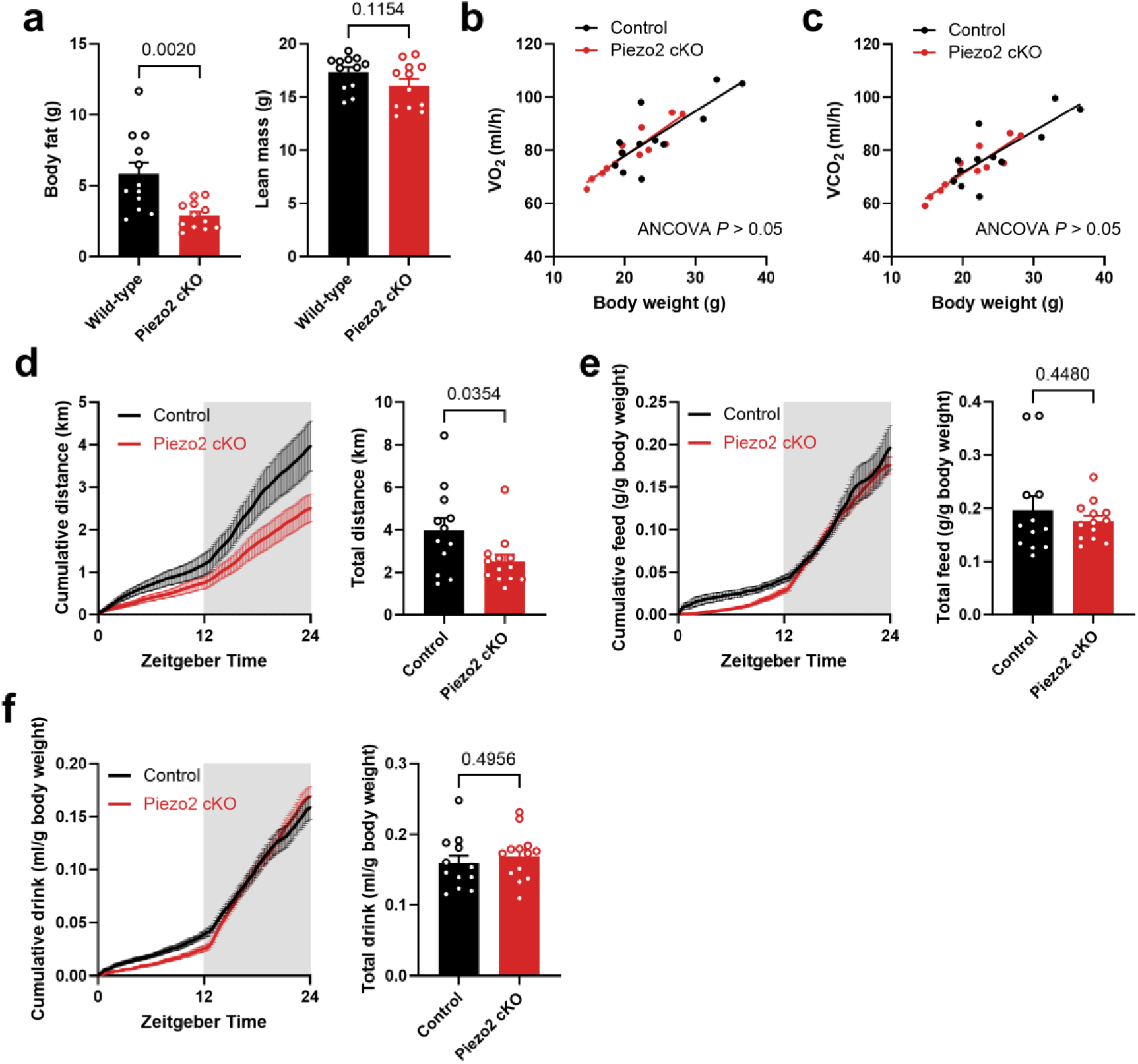
Body composition and calorimetric and behavioral assessment of *Piezo2* cKO mice. a) Body composition of *Piezo2* cKO mice and littermate controls (n=12 per group). b-f) Metabolic cage analysis provided data on b) oxygen consumption (VO_2_), c) carbon dioxide release (VCO_2_), d) mobility, e) food and f) water intake. Mice were allowed to habituate in the metabolic cages for three days, after that data were acquired. Data are the average of three consecutive days. Mice were singly housed at room temperature of around 22°C. Data in a and e-g were analyzed using an unpaired t-tests. Data in b,c were analyzed using ANCOVA with body weight as covariate. Data are mean ± SEM.

**Fig. S4.**
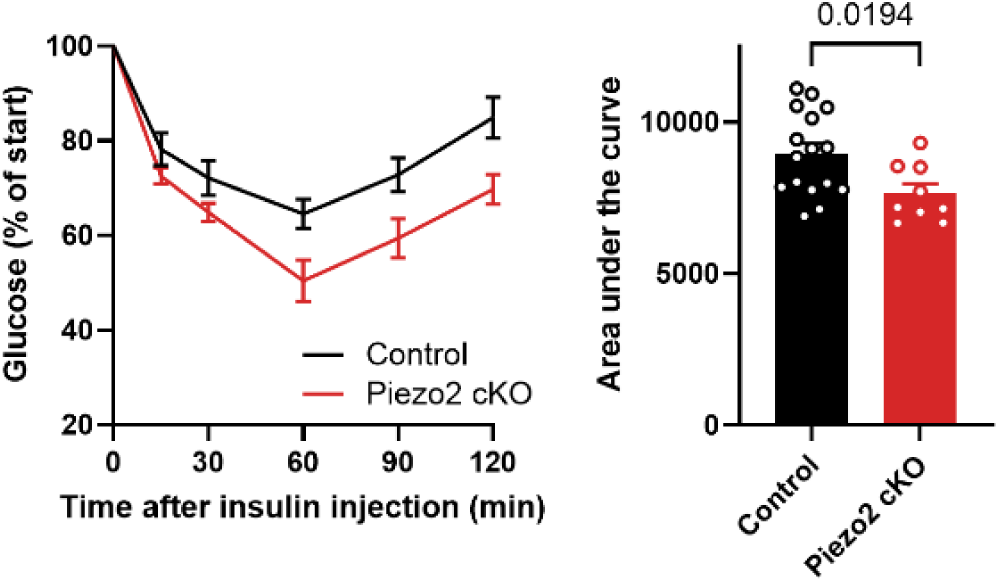
*Piezo2* cKO mice show increased insulin sensitivity. Insulin tolerance test (0.6U/kg body weight) was performed after 6 h fasting. Statistical analysis was performed using unpaired t-tests. n=9 *Piezo2* cKO mice, n=16 littermate controls. Data are mean ± SEM.

**Fig. S5.**
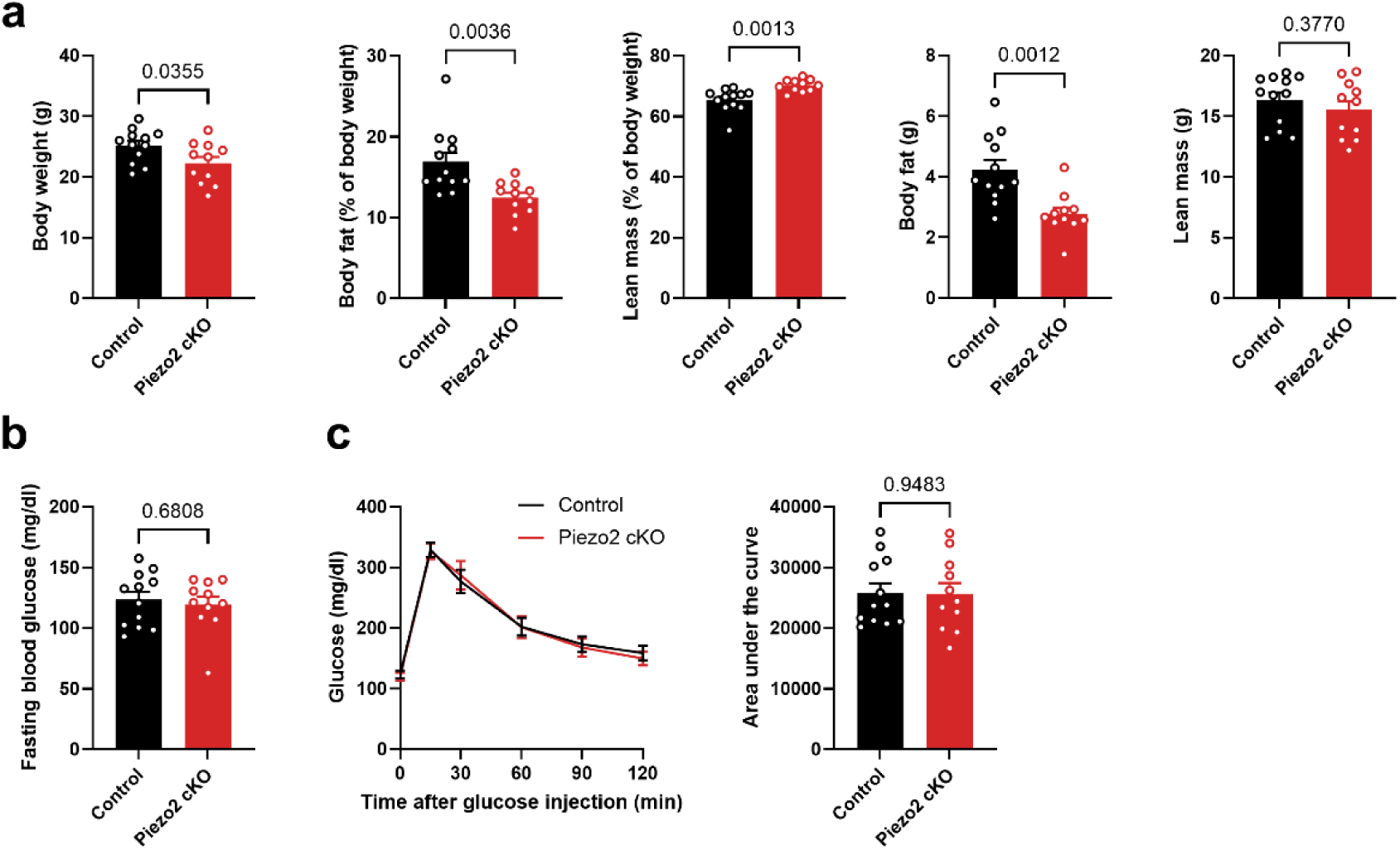
*Piezo2* cKO mice housed at thermoneutrality. a) Body composition analysis. b) Fasting blood glucose levels after 6 h fasting. c) Glucose tolerance test (2 g/kg body weight). Mice were housed at thermoneutrality (30°C) starting shortly after birth (p1-p3). *Piezo2* cKO mice and littermate controls were 3-6 months old at the timepoint of the experiments. Statistical analysis was performed using unpaired t-tests. n=12 *Piezo2* cKO mice, n=11 littermate controls. Data are mean ± SEM.

**Fig. S6.**
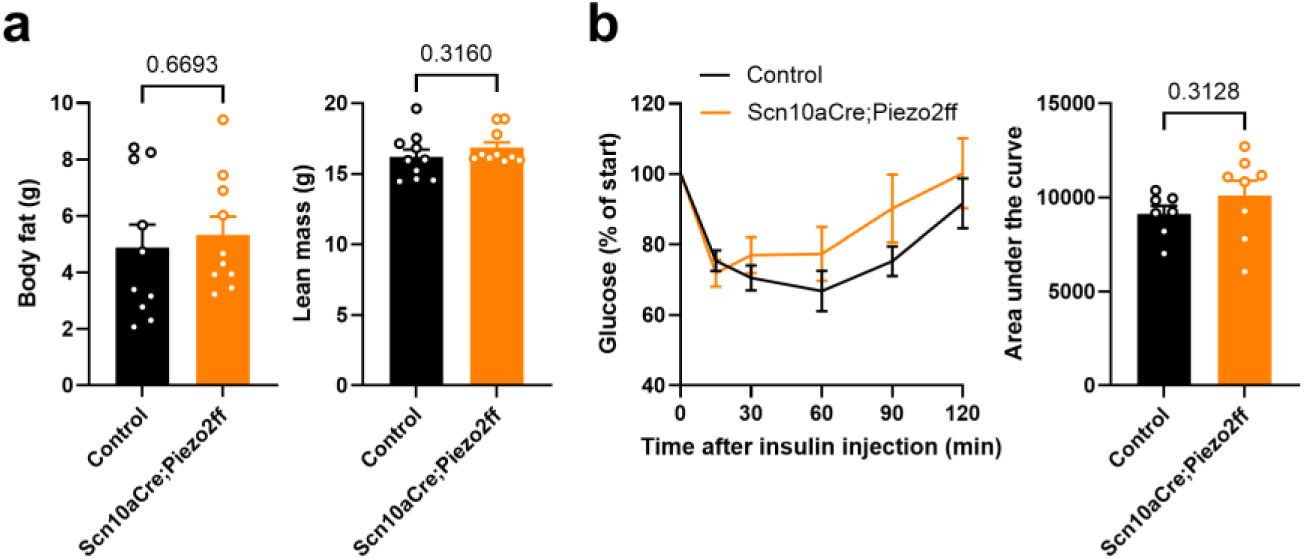
*Scn10aCre*;*Piezo2flox/flox* mice showed unchanged body composition and insulin sensitivity. a) Body composition of *Scn10a-Cre;Piezo2flox/flox* mice (n = 10) and littermate controls (n = 10). b) Insulin tolerance test (0.6 U/kg body weight) performed after 3 h fasting showed unaffected insulin sensitivity in *Scn10aCre;Piezo2flox/flox* mice (n = 8) compared to littermate controls (n = 7). Mice were kept at room temperature of around 22°C, and were 3-6 months old at the timepoint of the experiments. Statistical analysis was performed using unpaired t-tests. Data are mean ± SEM.

**Fig. S7.**
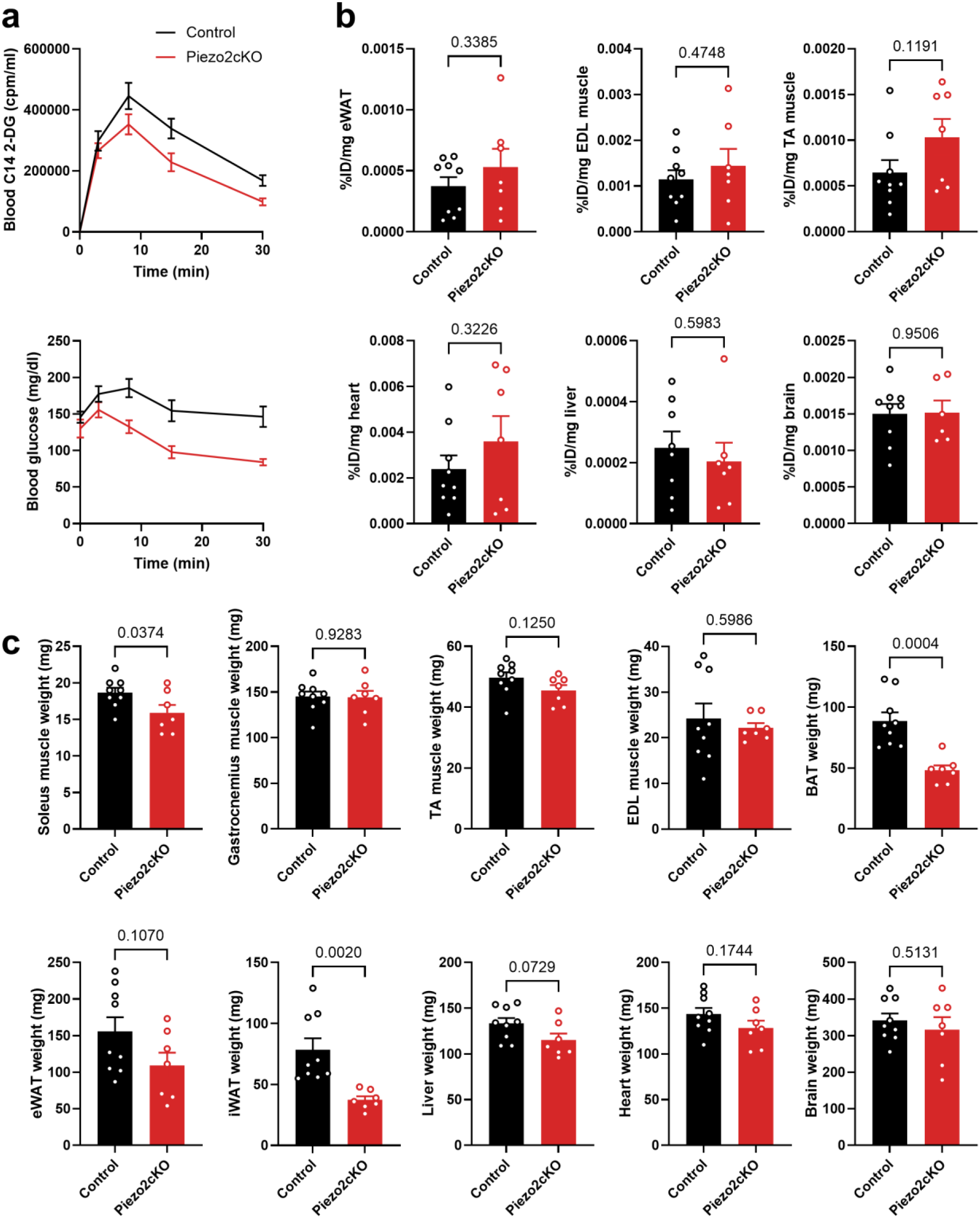
Tracing of radiolabeled 2-DG uptake. a) Blood levels of radiolabeled 2-DG and of glucose upon co-injection of 2-DG and insulin. b) 2-DG uptake in visceral epididymal white adipose tissue (eWAT), extensor digitorum longus (EDL) muscle, tibialis anterior (TA) muscle, heart, liver and brain. Data are presented as % of injected dose (ID) per mg tissue. c) Corresponding tissue weights. Statistical analysis was performed using unpaired t-tests. n=6-7 *Piezo2* cKO mice, n=9 littermate controls. Data are mean ± SEM.

**Fig. S8.**
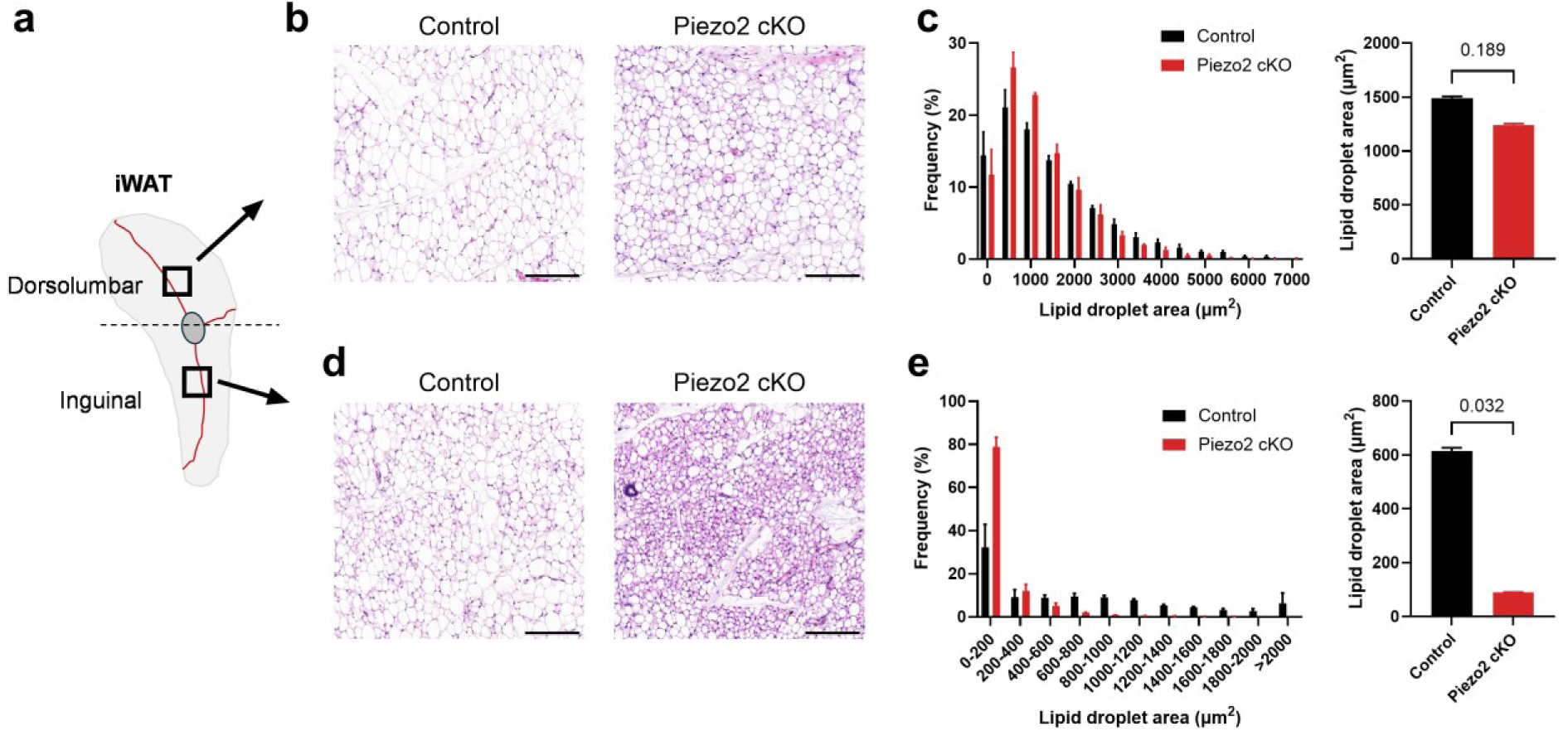
Reduced lipid droplet size in the iWAT of Piezo2 cKO mice particularly in the beiging region. a) Schematic illustrating the two analyzed areas: dorsolumbar and inguinal region. Beiging occurs more prominently in the inguinal region. b) H&E staining of mutant and control iWAT (dorsolumbar area); scale bar 200 µm. c) Quantification of lipid droplet area. d) H&E staining of mutant and control iWAT (inguinal area); scale bar 200 µm. e) Quantification of lipid droplet area. Statistical analysis was performed using a linear mixed model with the mouse ID as a random factor. n=3 *Piezo2* cKO mice, n=3 littermate controls. Data are mean ± SEM.

**Fig. S9.**
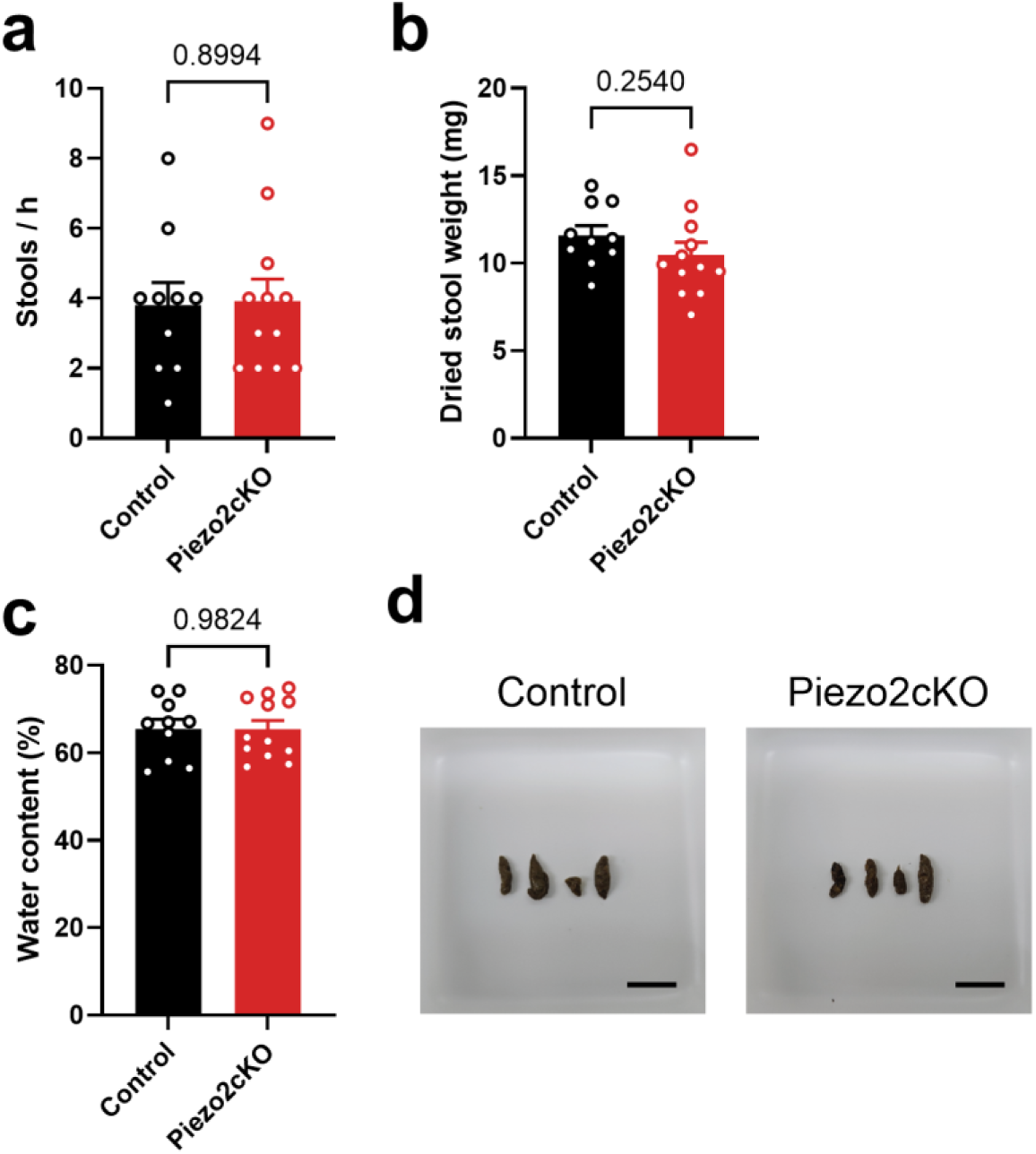
*Piezo2* cKO mice show no overt sign of gastrointestinal dysfunction. a) Number of stools collected within 1h from control and *Piezo2* cKO mice. b) Stools were dried and weighted individually and then averaged per mouse. c) Water content of the stools. d) Representative images of the stools from a control and a *Piezo2* cKO mouse. Scalebar, 1 cm. Statistical analysis was performed using unpaired t-tests. n=12 *Piezo2* cKO mice, n=10 littermate controls. Data are mean ± SEM.

**Figure S10:**
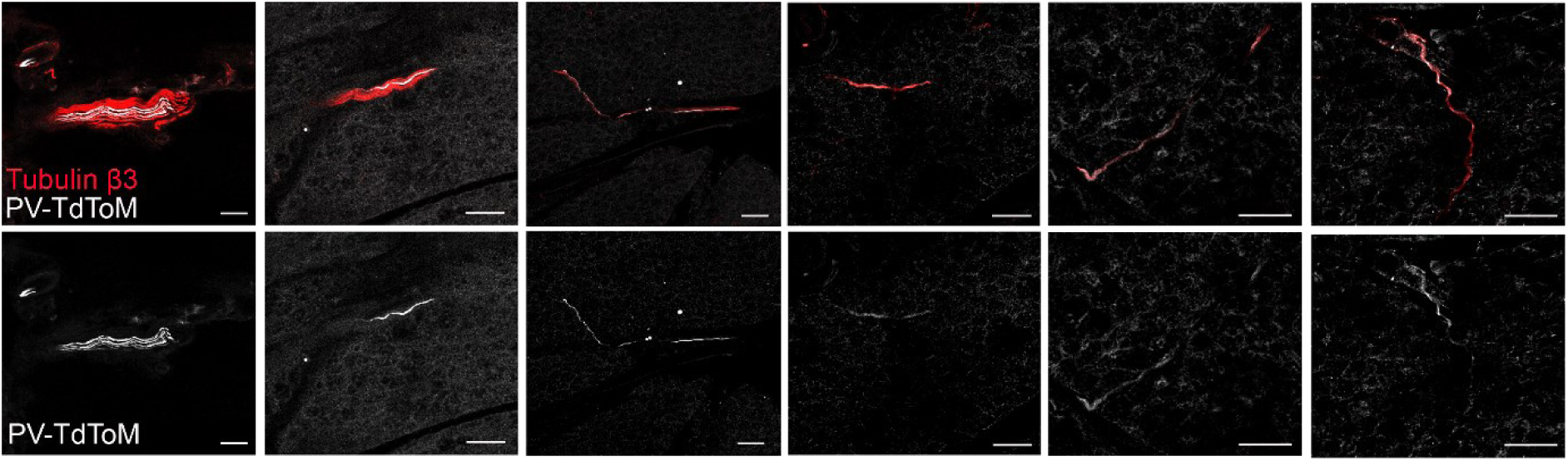
PV-positive innervation in iBAT. Representative images of PV-positive neurons in large nerve fibers (2 leftmost images) and thin nerve fibers (remaining images) inside the tissue. iBAT sections from PVCre;Tomato mice were stained for the pan-neuronal marker Tubulin β3 and TdTomato. Scale bar, 50 µm.

**Fig. S11.**
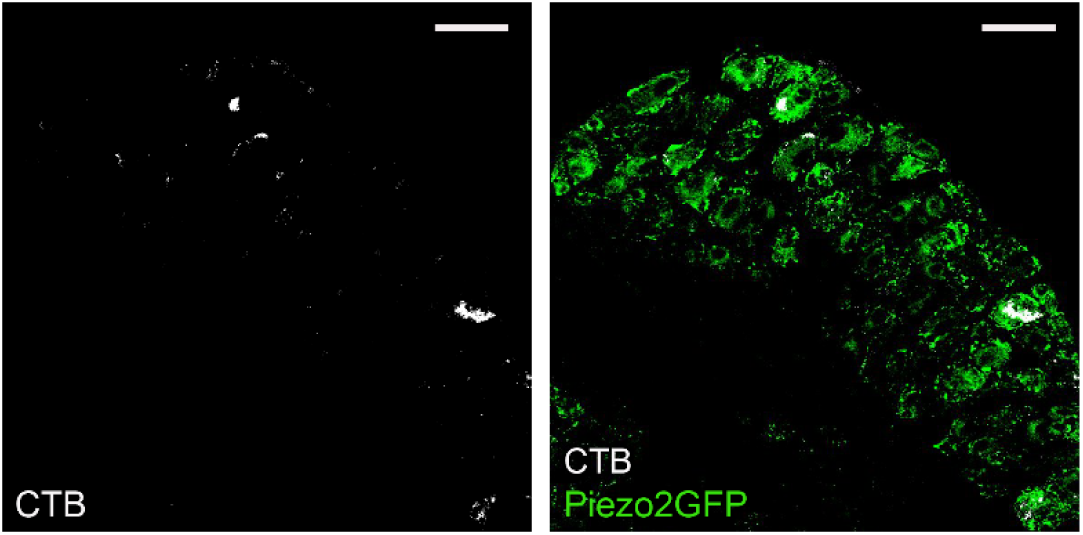
CTB labelling co-localizes with Piezo2GFP. Representative images of CTB labelling in T3 DRG of a Piezo2GFP mouse. Scalebar, 50 µm.

**Fig. S12.**
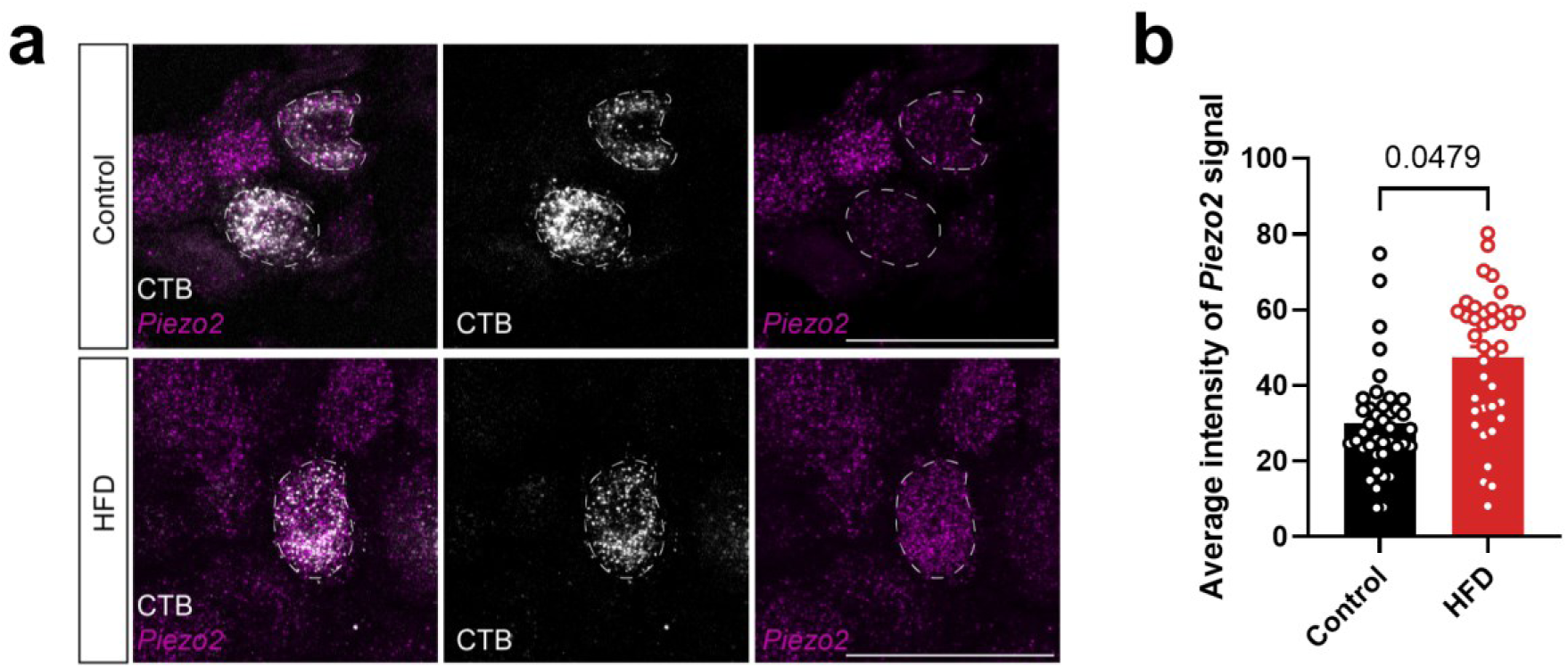
HCR smFISH for *Piezo2* in DRG sections with CTB labeling from iWAT of control and HFD-fed mice. a) Representative images of *Piezo2* smFISH and CTB labeling in T12-L1 DRG of control and HFD-fed mice (8 weeks of HFD). Scalebar, 50 µm. CTB was injected into iWAT. HFD mice receive 8 weeks of HFD (average body weight 40±2g), while control mice were maintained on standard chow diet (average body weight 33±1g). b) Quantification of *Piezo2* signal intensity in control mice (n = 3 mice with total of 40 CTB-positive cells) compared to HFD mice (n = 3 mice with total of 38 CTB-positive cells). Unpaired t-test was performed with the average signal for each mouse. Data are mean ± SEM.

**Table S1:**
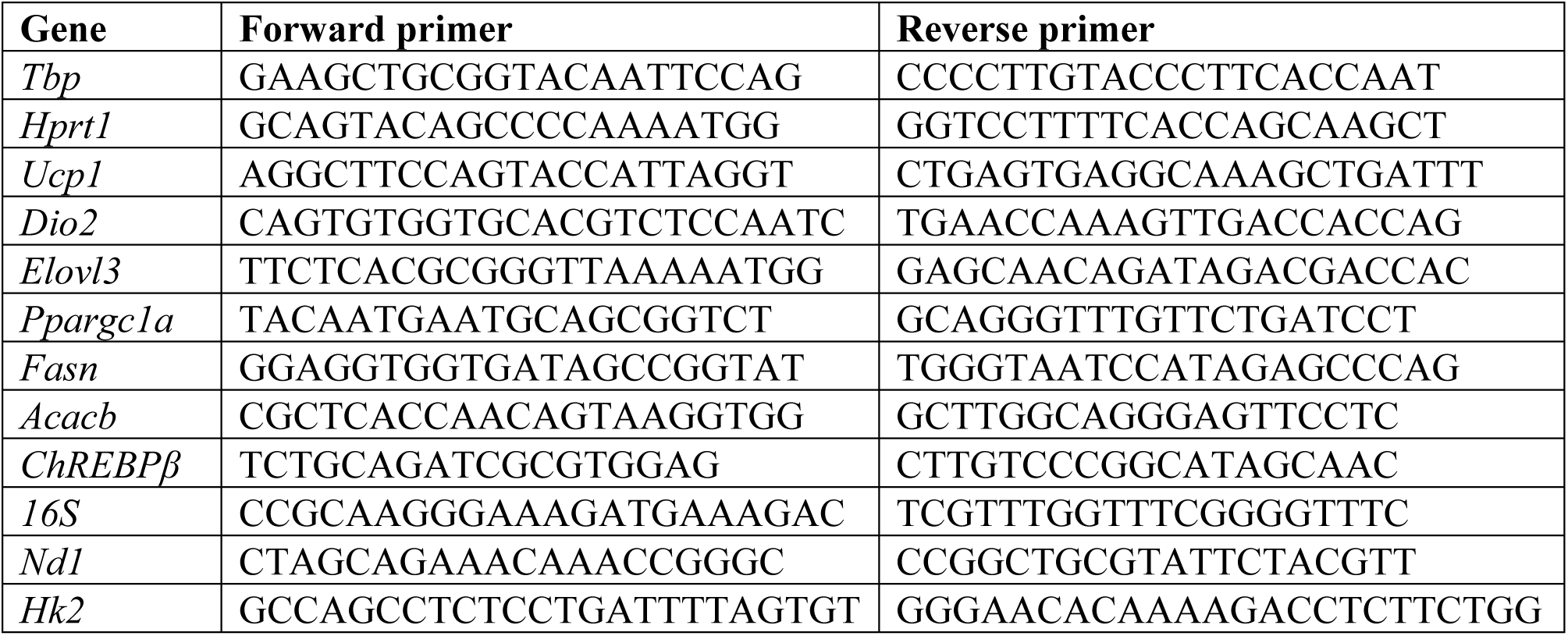
Forward and reverse primer sequences used for RT–qPCR.

